# Prominin-1 null Xenopus laevis develop subretinal drusenoid-like deposits, cone-rod dystrophy, and RPE atrophy

**DOI:** 10.1101/2024.06.03.597229

**Authors:** Brittany J. Carr, Dominic Skitsko, Jun Song, Zixuan Li, Myeong Jin Ju, Orson L. Moritz

## Abstract

Mutations in the *PROMININ-1* (*PROM1)* gene are associated with inherited, non-syndromic vision loss. Here, we used CRISPR/Cas9 to induce truncating *prom1*-null mutations in *Xenopus laevis* to create a disease model. We then tracked progression of retinal degeneration in these animals from the ages of 6 weeks to 3 years old. We found that retinal degeneration caused by *prom1*-null is age-dependent and likely involves death or damage to the retinal pigment epithelium (RPE) that precedes photoreceptor degeneration. As *prom1*-null frogs age, they develop large cellular debris deposits in the subretinal space and outer segment layer which resemble subretinal drusenoid deposits (SDD) in their location, histology, and representation in color fundus photography and optical coherence tomography (OCT). In older frogs, these SDD-like deposits accumulate in size and number, and they are present before retinal degeneration occurs. Evidence for an RPE origin of these deposits includes infiltration of pigment granules into the deposits, thinning of RPE as measured by OCT, and RPE disorganization as measured by histology and OCT. The appearance and accumulation of SDD-like deposits and RPE thinning and disorganization in our animal model suggests an underlying disease mechanism for *prom1*-null mediated blindness of death and dysfunction of the RPE preceding photoreceptor degeneration, instead of direct effects upon photoreceptor outer segment morphogenesis, as was previously hypothesized.

## INTRODUCTION

Inherited retinal degeneration is a heterogeneous set of disorders caused by mutations in genes important for retinal function that result in progressive vision loss and blindness. Advances in genetic screening have identified hundreds of genes and mutations potentially linked to inherited retinal degeneration. However, relatively little is known about the function of these genes, genotype-phenotype correlation for mutations, or underlying mechanisms by which mutation causes blindness. This includes retinal disorders linked to mutations in *PROMININ-1* (*PROM1*). *PROM1*-null mutations are associated with autosomal recessive cone-rod dystrophy, retinitis pigmentosa (RP), and macular degeneration (Maw et al., 2000; Permanyer et al., 2010; Pras et al., 2009; Zhang et al., 2007). The point-mutation *PROM ^R373C^* allele is linked to autosomal dominant Stargardt-like juvenile macular degeneration (Michaelides et al., 2010). Mouse studies concluded that retinal degeneration caused by *prom1* mutations was caused by defective outer segment disc morphogenesis (Yang et al., 2008; Zacchigna et al., 2009). Modelling *prom1*-null mutations in *Xenopus laevis*, however, has shown that although photoreceptor outer segments are dysmorphic, outer segment morphogenesis and visual function is maintained for at least 6 weeks (Carr et al., 2021). Here, we report that with age (≥ 6 months), small autofluorescent deposits appear in the outer segment layer between the inner segment/outer segment boundary and the retinal pigment epithelium (RPE) in *prom1*-null *X. laevis*. These deposits increase in size and number as animals continue aging and resemble human subretinal drusenoid deposits (SDD) (Zweifel et al., 2010). These findings call into question previous conclusions that disrupted outer segment morphogenesis is the direct cause of *prom1*-null associated retinal degeneration (Carr et al., 2021; Lu et al., 2019; Zacchigna et al., 2009). Rather, it appears that secondary toxic effects such as accumulation of these SDD-like deposits or dysfunction or death of RPE that occurs with age, may be instead responsible for photoreceptor death caused by *prom1*-null mutations.

Human SDD are also termed reticular pseudodrusen (RPD) and differ from conventional soft and hard drusen in their localization and composition (Wightman and Guymer, 2019). SDD localize to the subretinal space, between the retinal outer segments and the apical side of the RPE. Soft drusen are present external to the basal border of the RPE, between the RPE and Bruch’s membrane (Curcio, 2018). Oil Red O, a neutral lipid stain, colors soft drusen a vivid red but does not stain SDD, suggesting different lipid compositions and sources (Greferath et al., 2016; Oak et al., 2014). Human SDD contain unesterified cholesterol, apolipoprotein E (apoE), complement factor H, and vitronectin, while soft drusen contain apoB, apoA-1, and esterified cholesterol (Greferath et al., 2016; Rudolf et al., 2008; Sarks et al., 1988; Wightman and Guymer, 2019). However, the full protein and lipid composition of SDD is not known and there is controversy as to the presence of proteins like rod and cone opsins, which would indicate outer segment membrane components in SDD (Greferath et al., 2016; Rudolf et al., 2008).

The prevalence of SDD and their role in retinal degenerative disease has historically been unknown. This is due in part to the difficulty in detecting SDD with color fundus photography coupled with the lack of an animal model (Zarubina et al., 2016), (Fletcher et al., 2014). There is mounting evidence that SDD are present in the retinas of most people, irrespective of whether they have retinal degenerative disease (Zarubina et al., 2016). However, at some point, increased size or number of SDD tip the scales from a benign occurrence to a pathogenic trigger, increasing the risk for severe retinal degenerative disease, especially for dry (geographic atrophy) and wet (choroidal neovascularization, CNV) age-related macular degeneration (AMD) (Wightman and Guymer, 2019); the cause of this tipping point remains elusive. Here, we demonstrate that aging *prom1*-null *X. laevis* develop deposits of cellular debris that resemble SDD. Affected frogs show analogous effects to severe retinal degenerative disease and some hallmarks of dry AMD such as retinal atrophy, retinal pigment epithelium (RPE) migration and/or death, and the absence of CNV. These identify a potential novel target therapeutic pathway for *prom1*-null mutations: secondary toxic effects on the RPE. This also establishes *X. laevis* as a potential animal model for understanding some of the key hallmarks of dry AMD.

## METHODS

### Animal ethics statement, housing, & anesthesia

Animal use protocols were approved by the University of British Columbia (A18-0257/A18-0259) and University of Alberta (AUP00004203) Animal Care Committees and carried out in accordance with the Canadian Council on Animal Care. Animals were housed at 18°C under a 12-hour cyclic light schedule (900-1200 lux). For ERG and live imaging experiments, adult frogs were anesthetized in 0.3% tricaine buffered in 0.1x MMR (10 mM NaCl, 0.2 mM KCl, 0.1 mM MgCl_2_•6H_2_O, 0.2 mM CaCl_2_, 0.55 mM HEPES) until non-responsive (8-15 min). Adult frogs were not anesthetized more than twice in one week and were allowed to recover for a minimum of 48 hrs between anesthetic exposures.

Frogs used in this study are a combination of F0 and F1 animals that were created and characterized using the same methods as described previously (Carr et al., 2021). CRISPR-Cas9 genetic modification was performed using a single-guide RNA (sgRNA) that targeted exon 1 of the *prom1* L gene. Blood samples or heterozygous F1 embryos were used for genomic DNA extraction to confirm and characterize CRISPR-mediated indels by Sanger sequencing. Matings between animals that had confirmed indels by blood draw were performed to obtain F1 progeny.

### Immunofluorescence and Oil Red O stain

Whole eyes were fixed and prepared for immunohistochemistry as described previously(Carr et al., 2021). Tissues were mounted using medium prepared from Mowiol 4-88 (Mowiol mounting medium, 2006) (Millipore Sigma, St. Louis, MO, USA) and then visualized using a Zeiss LSM 800 microscope equipped with a 40× N.A. 1.2 water immersion objective, a Zeiss LSM 880 with Airyscan equipped with a 63×1.4 N.A. oil immersion objective, or a Leica DM600B epifluorescence microscope with a Leica K3M 6.3 MP CMOS camera. Micrographs represent maximum intensity projections of whole retinal sections (z = 0.3-0.5 µm/step) unless specified otherwise. Antibody sources, conjugates, and concentrations used in this study are listed in Supplementary Table S1.

Oil Red O (ORO; Millipore Sigma, St. Louis, MO, USA) was freshly prepared each time. 0.5% (w/v) ORO was dissolved in 100% isopropanol, filtered with a syringe filter (0.22 µm) and then mixed 3 parts ORO to 2 parts dH_2_O to prepare a 0.3% (w/v) staining solution diluted in 60% isopropanol. Tissue sections were washed 3 x 8 min in PBS to remove OCT, incubated in 60% isopropanol for 5 min, and then stained with 0.3% ORO for 15 min. Stained tissues were rinsed 2-5 times (until clear) with 60% isopropanol using a transfer pipette and then washed 3 x 8 min in PBS with gentle shaking. Tissues were then coverslipped using Mowiol mounting medium (Mowiol mounting medium, 2006) and imaged in brightfield using the Zeiss LSM 800 confocal microscope and objectives as described above. A Zeiss Axiocam 503 color camera was used to collect brightfield images.

### Transmission Electron Microscopy (TEM)

Detailed methods for TEM tissue preparation followed previously published protocols(Carr et al., 2021; Tam et al., 2015). Whole eyes were fixed in 4% paraformaldehyde + 1% glutaraldehyde diluted from 20% and 8% stock solutions, respectively (Electron Microscopy Sciences, Hatfield, PA, USA). Eyes were fixed at 4°C for ≥ 24 hrs and then infiltrated with 2.3 M sucrose, embedded in OCT (ThermoFisher, Waltham, MA, USA), and cryosectioned at 20 µm with single sections per slide. Optimally oriented sections were washed with 0.1 M sodium cacodylate and then stained with 1% osmium tetroxide for 30 min. After staining, sections were dehydrated in increasing concentrations of anhydrous ethanol (30%, 50%, 70%, 90%, 95%, each for 5 min, 100% 3 x 5 min) and then infiltrated with increasing concentrations of Eponate 12 resin (Ted Pella Inc., Redding, CA, USA) diluted with anhydrous acetone (35%, 50%, 70%, each for 40 min, 100% 2 x 40 min). Beem capsules with the ends trimmed off were placed over the fully infiltrated sections and then the capsule was filled with resin and allowed to polymerize overnight. Ultrathin silver-grey were cut with a diamond knife and collected on Formvar-coated nickel slot grids (Electron Microscopy Sciences, Hatfield, PA, USA). Sections were stained with saturated aqueous uranyl acetate and Venable and Coggeshall’s lead citrate (Venable and Coggeshall, 1965). Imaging was performed with a JEOL 2100 at 200 kv.

### Optical Coherence Tomography, Fundus Photography, and Fluorescein Angiography

Infrared optical coherence tomography (OCT) and color fundus images were taken using a Micron IV system with the mouse-specific objective lens (Phoenix Technology Group, Pleasanton, CA). Brightfield color fundus images had a field of view of 1.8 mm. The infrared OCT spectrometer covered 750-930 nm and the full line scan size was 1.6 mm in X and Y. Micron IV OCT images were created from 15 averaged segments. For fluorescein angiography, 25% fluorescein sodium (Millipore Sigma, St. Louis, MO, USA) diluted in sterile dH_2_O was injected into the dorsal lymph sac (2 µL/g of frog). The initial concentration was based on previous studies in mouse (Chang, 2013) and then optimized empirically for this study. The resulting retinal blood vessel fluorescence was imaged using the Micron IV colour fundus camera at its peak intensity, 20-30s after injection.

Spectral-domain optical coherence tomography (SD-OCT) was performed with a custom-built system tailored for small animal retinal imaging (Wahl et al., 2019). The system utilized a superluminescent diode laser source with the center wavelength of 840 nm and a full-width-half-maximum bandwidth of 100 nm yielding axial resolution of approximately 3.1 µm. The detection of the OCT signal was performed by two separate custom-built spectrometers whose maximum A-scan rate was 250 kHz and numerically calibrated to acquire identical back-scattering signals from the subject with an opposite polarity (Miao et al., 2022). The optical signals acquired from the spectrometers were processed to extract melanin granule-specific contrast from the subject by estimating a noise-suppressed degree of polarization uniformity (DOPU) whose processing algorithm adhere to a protocol developed previously (Hsu et al., 2020; Makita et al., 2014). Each processed multi-contrast degree of polarization uniformity (DOPU) retinal OCT image presented in this study was constructed with 450 A-scans/B-scan and 500 B-scans/volume.

### Electroretinography

Electroretinograms (ERGs) were recorded using electrodes connected to a model 1800 AC amplifier and head stage (AM Systems, Sequim, WA) and an Espion Ganzfeld stimulator (ColorDome) and recording unit (Diagnosys LLC, Lowell, MA) as described previously (Vent-Schmidt et al., 2017) with minor modifications as follows: The ground, a flat gold EEG electrode, was glued into a 100 mm petri dish instead of a 60 mm petri dish. The chin of the adult frogs was placed on the ground electrode. To accommodate the rest of the body, 1/3 of the sides of the 100 mm petri dish were cut off, smoothed with sandpaper, and then lined with soft weather stripping to prevent scratching the belly of the frogs and to keep the frogs from sliding out of the dish during ERG recordings. Corneal recordings were made from a chlorinated silver wire electrode set in a pulled glass micropipette and mounted into an electrode pin holder (A-M Systems, Sequim, WA). The entire electrode assembly was mounted in a manual micromanipulator (US-3F, Narishige International, Amityville, NY, USA). Adult frogs were anesthetized in 0.1% Tricaine in 0.1x MMR until just unresponsive, as deep anesthesia can negatively impact the strength of the ERG response (Nair et al., 2011). Scotopic ERGs were recorded in animals that had been dark-adapted overnight and prepared under dim red light; recorded stimuli were the average of 5 trials. Photopic ERGs were recorded in animals that had been exposed to a normal light cycle and prepared under regular lab lighting (∼350 lux); recorded stimuli were the average of 10 trials. For scotopic and photopic ERG, the A-wave amplitude was measured as the minimum value (trough) between 50-150 ms post-flash stimulus. The B-wave amplitude was measured as the value from the a-wave trough to the b-wave peak, which was defined as the maximum value measured between 100-250 ms post-flash stimulus. Flicker amplitude was measured as the average of the values from trough to peak for 5 cycles.

### Image analysis for prom1 immunoreactivity and shed outer segment tips

Prom1 immunolabeling signal intensity was measured in FIJI using a macro created by Dr. Elizabeth Kugler (EKugler_11082021.ijm). Stitched micrographs were acquired using the same image settings and then raw data files were imported into FIJI. Thresholding was applied to the WGA channel, which facilitated automatic drawing of outer segment ROIs. Thresholded ROIs were used to measure summed pixel intensity (Raw Integrated Density) in both the WGA and prom1 channels, and the prom1 signal was then normalized to the WGA signal for each ROI using the formula (prom1 ÷ WGA x 100). All normalized ROI calculations were averaged for each age group and plotted in GraphPad Prism. Because data were not normally distributed, a Mann-Whitney U test was used to compare overall prom1/WGA normalized signal intensity.

To quantify outer segment disc shedding, tadpoles were raised completely in the dark for 2 weeks. On the morning of day 15 (around 10:00 am), tadpoles were exposed to 10-15 min of laboratory light (∼500 lux), after which the animals were euthanized, the eyes immediately fixed, and then the tissues were processed for immunohistochemistry as described above. To count the size and number of shed tips, regions of interest were manually drawn around visible shed outer segment tips in fluorescence micrographs of the central retinal representing a field of view of 160 µm x 160 µm. The ROI number and area was calculated using the measure function in FIJI (Schindelin et al., 2012).

### Statistical Analysis

ERG waveforms were visualized using Excel (Microsoft, Redmond, WA) and analyzed by measuring and plotting the A-wave and B-wave peak amplitudes, and then fitting the resulting curves using non-linear regression analysis. Genotype effects, light intensity effects, and interaction effects were analysed by two-way ANOVA with Sidak’s post hoc test. Statistical analysis and graphing were performed using GraphPad Prism (V10.2.2; San Diego, CA, USA). Micrographs were analyzed using Photoshop (Adobe, San Jose, CAL), Affinity Photo (Serif, West Bridgford, UK) and FIJI (Schindelin et al., 2012).

## RESULTS

### Aging *prom1*-null animals develop large deposits of cellular debris in the outer segment layer that resemble SDD

Loss of prom1 protein resulted in dysmorphic rod and cone outer segments. Rod outer segments comprised large overgrown whorls of membranes within the plasma membrane, whereas the cone outer segments were elongated and fragmented, with cone opsin positive material dotting the outside of rod outer segments (Supplementary Fig. S1). Although the photoreceptors were severely dysmorphic, *prom1*-null retinas did not degenerate quickly. Instead, small puncta that labeled heavily with Hoechst nuclear stain appeared in the outer segment layer by 6 weeks (Fig. 1A-F).

**Figure 1.**
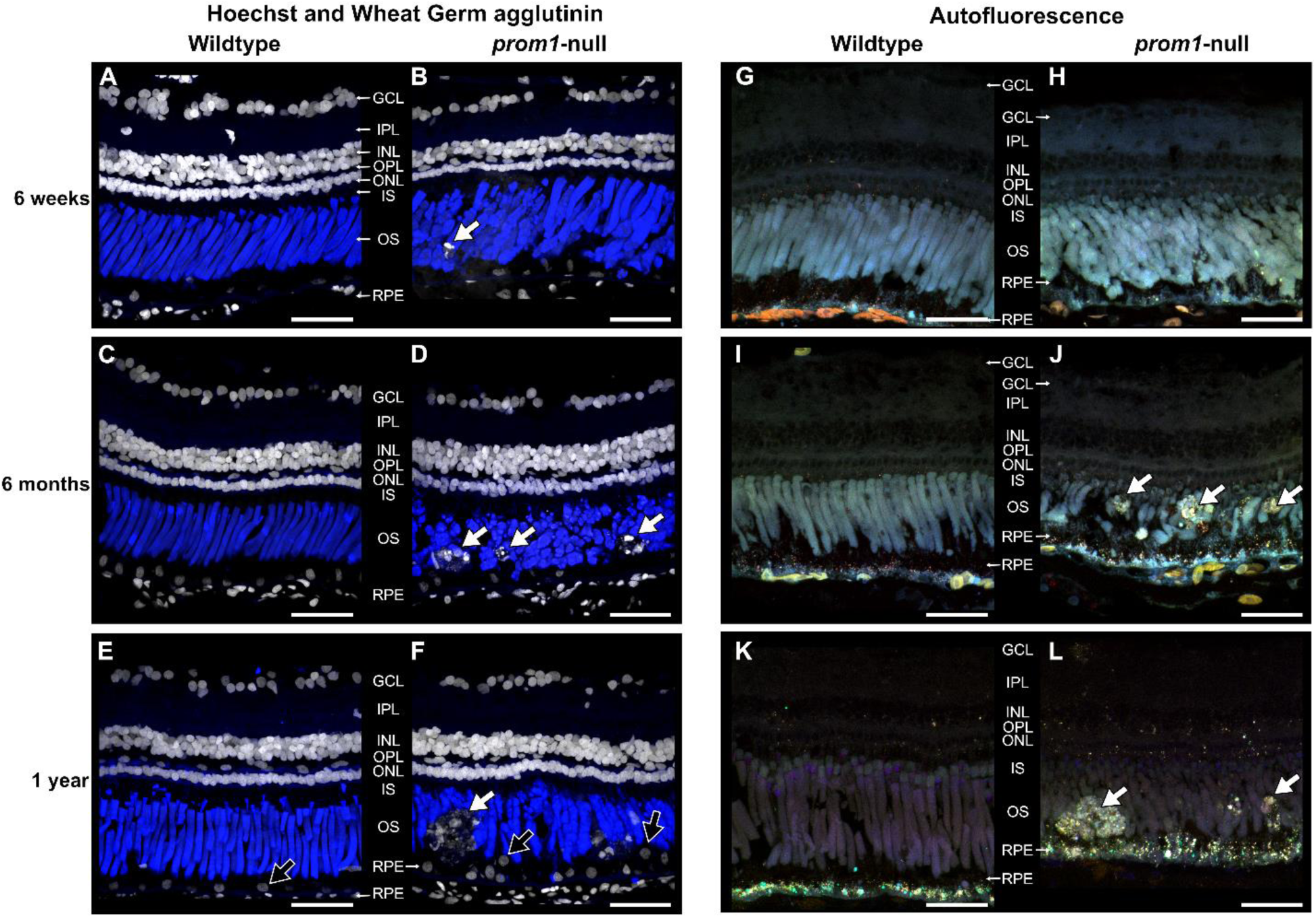
Representative histology of wildtype and F0 *prom1*-null retinas in animals aged 6 weeks to 1 year examined with Hoechst and WGA labeling. (A-F) and 4-channel label-less autofluorescence combining 405 nm, 488 nm, 555 nm, and 647 nm excitation wavelengths (G-L). Between 6-weeks and 1-year of age, deposits of cellular debris in the outer segment layer accumulate and increase significantly in size (white arrows). These deposits are strongly labelled with Hoechst dye (white; A-F) and are not present in wildtype control animals. Hoechst dye also demonstrates the progressive disorganization of RPE nuclei (F, black arrows). When examined using autofluorescence (G-L), the deposits fluoresce strongly at different excitation spectra than the surrounding outer segment material, and there is an increase in autofluorescent puncta in the RPE layer of prom1-null animals (H, J, L) compared to the wildtype controls (G, I, K). Autofluorescence images are color-coded to correspond to the detected emission spectra: cyan (405 nm), green (488 nm), orange (555 nm), and red (647 nm). Scale bars = 50 µm. *Labels (A-F):* Blue = wheat germ agglutinin (WGA); White = Hoechst nuclear dye. *Abbreviations:* GCL, ganglion cell layer; IPL, inner plexiform layer; INL, inner nuclear layer; OPL, outer plexiform layer; ONL, outer nuclear layer; IS, inner segments; OS, outer segments; RPE, retinal pigment epithelium. *Number of animals:* (A, n = 10; B, n = 13; C, n = 5; D, n = 5; E, n = 3; F, n = 6; G, n = 4; H, n = 4; I, n = 3; J, n = 5; K, n = 3; L, n = 6).

By 6 months, these puncta coalesced into larger deposits of cellular debris that resembled human SDD in localization, and by one year, very large deposits (35-60 µm) were present (Fig. 1). These large SDD-like deposits autofluoresced at wavelengths distinct from the surrounding outer segment material when viewed with label-less 4-channel confocal microscopy and there was an concurrent increase in lipofuscin-like autofluorescence in the RPE of *prom1*-null animals compared to wildtype animals (Fig 1G-L). In addition to the presence of large SDD-like deposits in the outer segment layer, organizational defects in the RPE were visible by displaced nuclei (Fig. 1F, black arrows) and increased RPE autofluorescence in *prom1*-null mutants at one year of age (Fig. 1L). Interestingly, we observed during our investigations that prom1 immunoreactivity decreased in the central retina of wildtype animals with full length rod outer segments (42 dpf) when compared to much younger animals, which have outer segments that are actively elongating in addition to maintaining daily disc synthesis (14 dpf). Prom1 immunoreactivity remained strong in the periphery of adult eyes, where outer segments were younger (Supplementary Fig. S2).

To investigate further similarities between these *X. laevis* SDD-like deposits and human SDD, labeling was performed with Oil Red O and BODIPY, which labels neural lipids but does not label human SDD. Our frog SDD-like deposits did not label strongly with Oil Red O or BODIPY (Fig. 2), indicating a lack of neutral lipids such as cholesterol esters and triacylglycerol (Bharati et al., 2022; Elle et al., 2010; Ramírez-Zacarías et al., 1992) and mimicking Oil Red O labeling results in human SDD. Brightfield and red autofluorescence imaging also revealed that the RPE in *prom1*-null animals was fragmented, extended much further into the outer segment layer, and infiltrated into the SDD-like deposits when compared to wildtype animals (Fig. 2).

**Figure 2.**
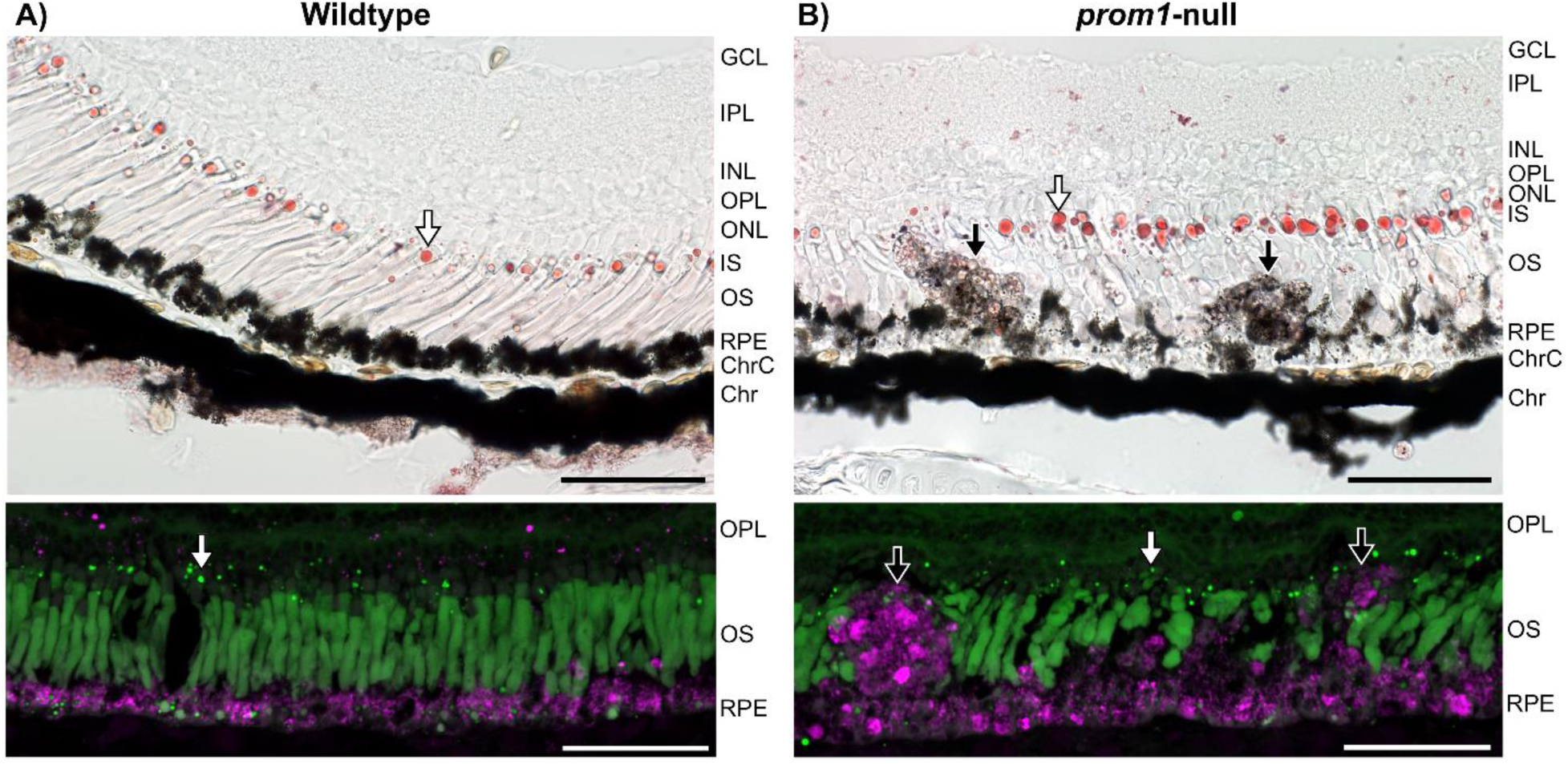
Representative images of Oil Red O and BODPIPY staining in 1-year old wildtype. (A) and *prom1*-null (B) retinas. Both Oil Red O and BODIPY label the oil droplets in the cone photoreceptors (white arrows), but neither labels the SDD-like deposits in the *prom1*-null retinas strongly (black arrows). The RPE is disrupted in *prom1*-null retinas; pigment granules infiltrate into the SDD-like deposits and migrate further into the outer segment layer than in wildtype animals. Red autofluorescence, is detected in the RPE of wildtype animals and in the RPE and large deposits in *prom1*-null animals (magenta, bottom panels). The outer segment layer is thinner in *prom1*-null animals compared to the wildtype counterparts. Scale bars = 50 µm. *Number of animals:* Wildtype, n = 5; *prom1*-null, n = 9. *Abbreviations:* GCL, ganglion cell layer; IPL, inner plexiform layer; INL, inner nuclear layer; OPL, outer plexiform layer; ONL, outer nuclear layer; IS, inner segments; OS, outer segments; RPE, retinal pigment epithelium; ChrC, choriocapillaris; Chr, choroid.

The SDD-like deposits in *prom1*-null animals did not contain significant amounts of rhodopsin or wheat germ agglutinin positive labeling, but did contain a small amount of cone opsin-positive material that showed up as a honeycomb-like shape within the deposits (Fig. 3A). Large deposits made up of multiple labeled nuclei did not contain RPE65 labeling, but some single nuclei displaced into the outer segment layer were surrounded by a ring of strong RPE65-positive immunoreactivity (Fig. 3B). The relative level of vimentin-positive signal was higher in the SDD-like deposits and in the RPE surrounding these deposits in *prom1*-null animals when compared to wildtype counterparts (Fig. 3C).

**Figure 3.**
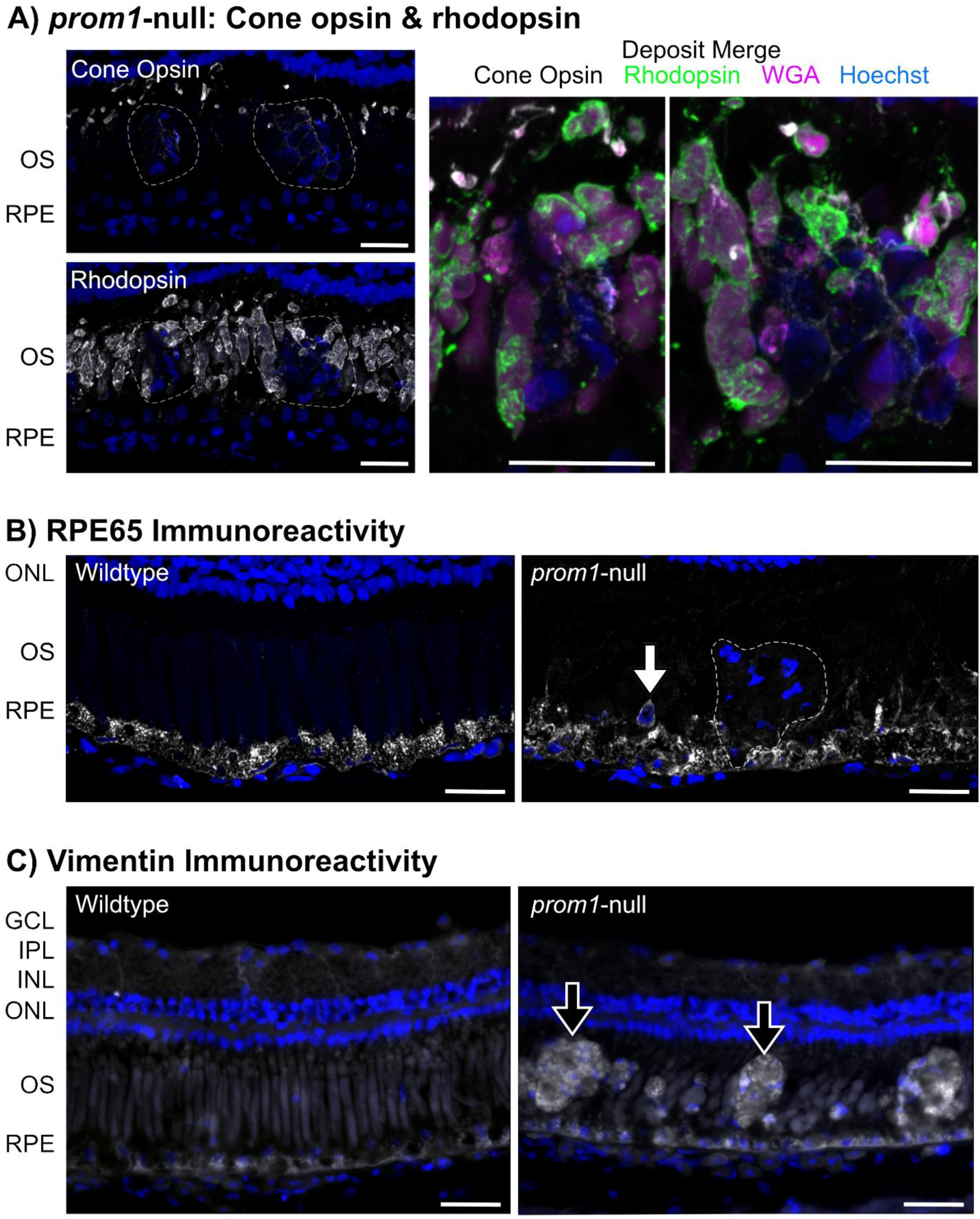
Labeling of SDD like deposits with markers for. (A) outer segments, (B) RPE, and (C) vimentin. SDD-like deposits do not show appreciable fluorescent signal for outer segment markers WGA, rhodopsin, or cone opsin (A, merge). Cone opsin was not detected in the deposits, though cone opsin positive-label surrounded the deposits in a honeycomb-like pattern (A, cone opsin, merge). Large deposits containing multiple nuclei are negative for RPE65 (B, dotted line), but single nuclei displaced from the RPE into the outer segment layer are surrounded by a ring of strongly RPE65-positive immunoreactivity (B; white arrow). Vimentin-positive immunoreactivity is increased in the RPE of *prom1*-null animals as well as the SDD-like deposits (black arrows), but stays relatively consistent in the cytoskeleton of the Müller glia in the inner plexiform layer (C). Scale bars = 25 µm. *Abbreviations:* GCL, ganglion cell layer; IPL, inner plexiform layer; INL, inner nuclear layer; ONL, outer nuclear layer; OS, outer segments; RPE, retinal pigment epithelium. Numbers of animals: Wildtype, n = 3-6; *prom1*-null, n = 3-5.

SDD-like deposits in *prom1*-null frogs were visible using color fundus photography and infrared optical coherence tomography (OCT) (Fig. 4). In F1 animals, the deposits showed up in color fundus photography as blue-white or yellow-white lesions that formed a consistent mosaic pattern throughout the entire retina (Fig. 4B,C). This finding is consistent with the *X. laevis* retinal mosaic of nearly equally spaced rods (53%) and cones (47%) and lack of a macula or visual streak (Gábriel and Wilhelm, 2001). Some frogs also exhibited widow defects which revealed the choroidal vasculature in addition to lesions (Fig. 4C). OCT confirmed that these deposits were located in the outer segment layer, between the photoreceptor inner segments and the apical side of the RPE, and penetration of the OCT beam through the RPE into the choroid (Fig. 4B,C). A detailed map of retinal layers visible with OCT in *X. laevis*, is shown in Supplementary Fig. S3. OCT segmentation and measurement indicated that *prom1*-null retinas were, on average, 13 ± 2.9 µm thinner than wildtype frogs (p < 0.05). The largest contributors to decreased retinal thickness were the outer segment layer (–3.5 ± 1.6 µm, n.s.) and the RPE (–7.9 ± 1.0 µm, p < 0.01; Supplementary Fig. S4).

**Figure 4.**
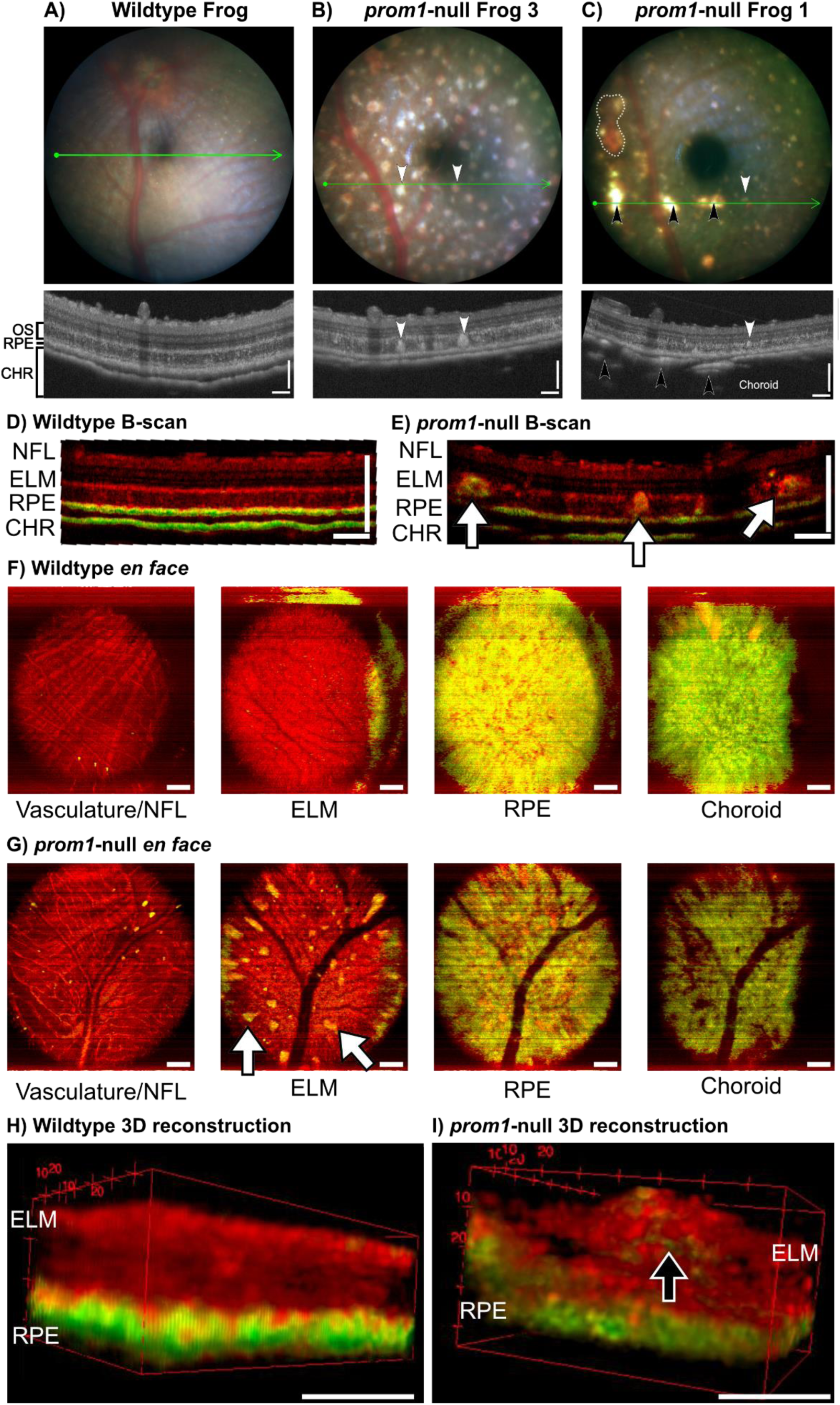
A-C) Representative images *prom1*-null F1 frogs compared to wildtype animals in color fundus photography and infrared OCT. F1 frogs have SDD-like deposits of cellular debris in the outer segment layer (white arrowheads), but one phenotype has a large number of these deposits and no window defects or penetration of the OCT beam through the dense RPE layer and into the choroid (B), while the other has fewer or no SDD-like deposits and large window defects that reveal the underlying choroid and choroidal vasculature (black arrowheads, C). Black arrows mark blood vessels on the vitreal side of the retina and the white arrow marks a large choroidal vessel that can be seen through the RPE in both the color fundus photo and the OCT scan. Fundus field of view = 1.8 mm. OCT Scale bars = 100 µm. D-I) Comparison of a wildtype animal and a 3 year old *prom1*-null adult frog with significant deposits using DOPU OCT in B-scan (D,E), *en face* (F,G), and 3D reconstructions (H,I). Deposits in *prom1*-null frogs have a significantly altered DOPU signal and are easily detectible as bright yellow or green spots in B-scan and *en face* orientations (E,G; white arrows). When projected in 3D, DOPU signal is not uniform within the deposits. Instead, green signal that indicates altered polarization caused by RPE pigment granules or other light-reflecting structures exists as a core within the deposit (I, black arrow). DOPU OCT detects loss of structural integrity of the ELM as evidenced by the projection of the large deposits through this layer, and loss of a strong band of red signal in the B-scan (E), *en face* (G), and 3D reconstructions (I). Evidence of RPE integrity breakdown is shown by breakdown of bright green bands in the B-Scan (E) and patchiness in the yellow *en face* image (G). The large black vessels in the *prom1*-null animal en face ELM, RPE, and choroid images (G) is not an anatomical defect, but is instead caused by shadowing from the vitreal vessels blocking the DOPU OCT signal. Scale bars = 200 µm. *Abbreviations:* NFL, nerve fibre layer; ELM, external limiting membrane; RPE, retinal pigment epithelium; CHR, choroid. *Number of animals:* wildtype = 6, *prom1*-null = 9.

SDD-like deposits were also visible using DOPU OCT as yellow-green lesions, indicating non-uniform polarization compared to surrounding retinal tissues (Fig. 4). Wildtype animals had uniform bright yellow-green external limiting membrane (ELM), RPE, and choroidal layers when examined with DOPU (Fig. 4D,F), while *prom1*-null mutants with severe disease phenotypes and large numbers of deposits exhibited more patchiness in these layers (Fig. 4E,G). The largest of the SDD-like deposits had a “core” of non-uniform polarization surrounded by uniform polarization (Fig. 4E, I) and they extended past the external limiting membrane, into the inner nuclear layer (Fig. 4E,G).

In addition to patchiness and breakdown of the RPE as observed with light microscopy (Fig. 2, Fig. 3) and OCT (Fig. 4), TEM of the RPE and outer segment tips of animals aged 8 months with moderate *prom1*-null disease revealed accumulations of complex, disorganized, electron dense deposits that contained putative membranes, lysosomes, vacuoles, and multivesicular bodies (Fig. 5). These bodies were located in or near the RPE and could represent lipofuscins or another type of indigestible waste by-product (Fig. 5A,B) as well as outer segment tip shedding defects (Fig. 5C). To investigate the potential for defective disc shedding in *prom1*-null mutants further, we quantified the size and number of shed outer segment tips released shortly after dark adaptation. We observed that, as early as two weeks post-fertilization, there were differences in outer segment shedding between *prom1*-null and wildtype animals. Shed membranes from *prom1*-null outer segment tips were smaller (4.2 ± 0.5 µm^2^) and more numerous (27.7 ± 5.1) than those shed by wildtype animals reared under the same conditions (9.1 ± 2.1 µm^2^, 11.0 ± 4.2, p < 0.05; Supplementary Fig. S5).

**Figure 5.**
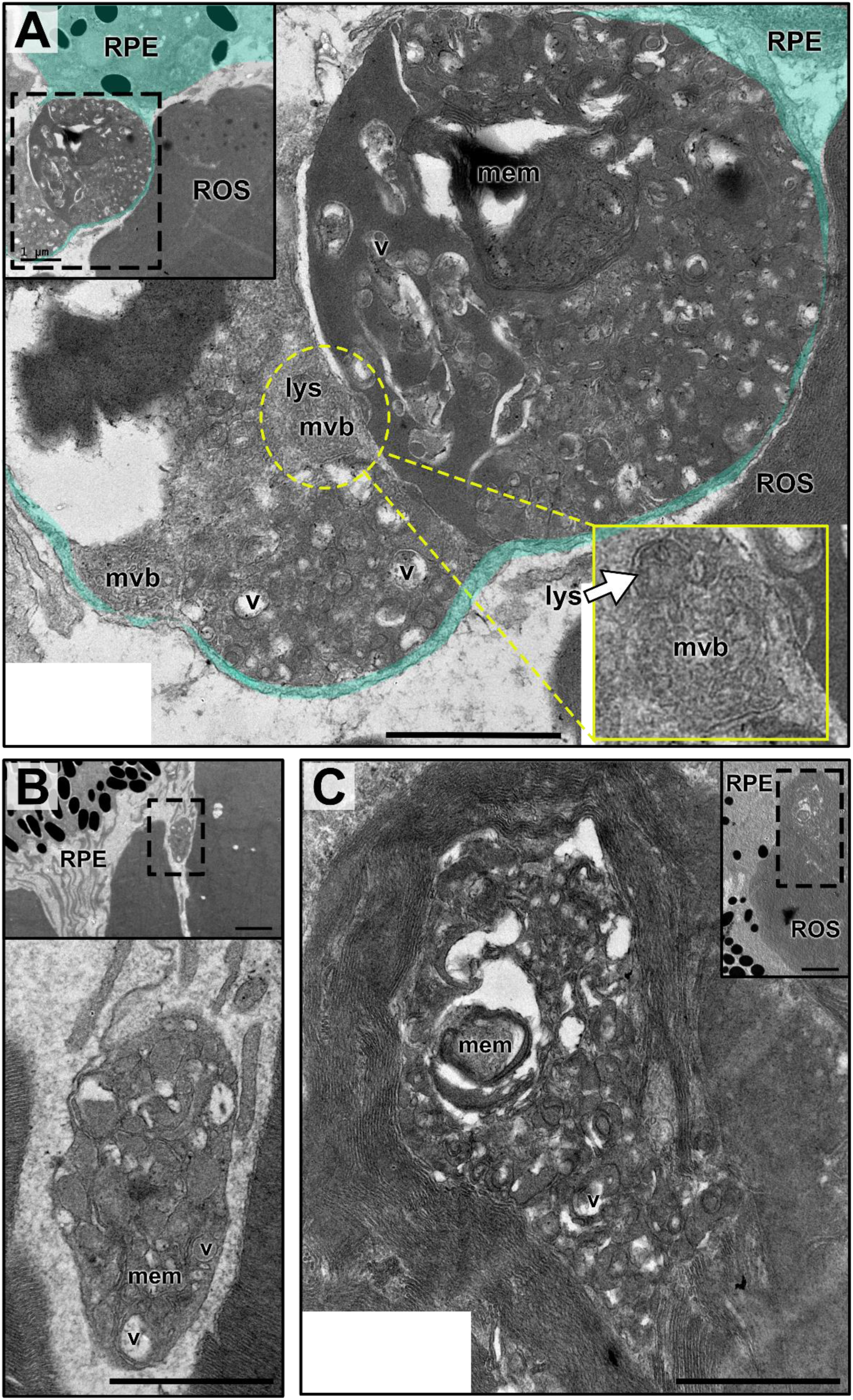
TEM micrographs of complex, lipofuscin-like deposits in the RPE of *prom1*-null mutants with a moderate disease phenotype (age 8 months). These deposits are localized in (A,B) or near (C) the RPE and contain electron dense material, membranes (mem), multivesicular bodies (mvb), lysosomes (lys), and numerous vacuoles (v). The yellow inset shows a higher magnification of a putative lysosome fusing to a disorganized multivesicular body in a large deposit located within the RPE. Scale bar = 0.5 µm, inset scale bar = 1 µm. *Abbreviations*: ROS, rod outer segment; RPE, retinal pigment epithelium.

The scotopic ERG response of animals aged 6 weeks was slightly impaired, but not significantly, and there was no progressive impairment as animals aged to 1 and 2 years old (Fig. 6). There was no significant difference in A-wave amplitude between wildtype and *prom1*-null animals at any time point (Fig. 6B,D). The B-wave amplitude was slightly impaired in *prom1*-null animals at 1 and 2 years old compared to 6 weeks, however this difference was small, and only statistically significant for 250 cd s/m^2^ between 6 weeks and 1 year of age (Fig. 6C,E; p < 0.05). The photopic ERG was significantly impaired at 6 weeks, 1 year, and 2 years of age but it did not disappear completely, not did this impairment continue to progress significantly between 6 weeks and 2 years old for flash ERGs (Fig. 7). The photopic A-wave was significantly impaired at all time points in *prom1*-null mutants compared to wildtype animals (Fig. 7B, Table 2), but the difference between wildtype and *prom1*-animals remained the same up to two years (Fig. 7E). This was also true for the photopic flash B-wave; it was significantly impaired for *prom1*-null animals at all ages (Fig. 7C), but the impairment of the B-wave did not progress up to two years (Fig. 7F). The only significant progressive impairment observed between 6 weeks and 2 years of age was high-intensity photopic flicker (Fig. 7G). 2-year-old frogs had an increased impairment in flicker response at 25 (p < 0.01) and 75 cd s/m2 (p < 0.01) compared to 1 year and 6 week old frogs, respectively.

**Figure 6.**
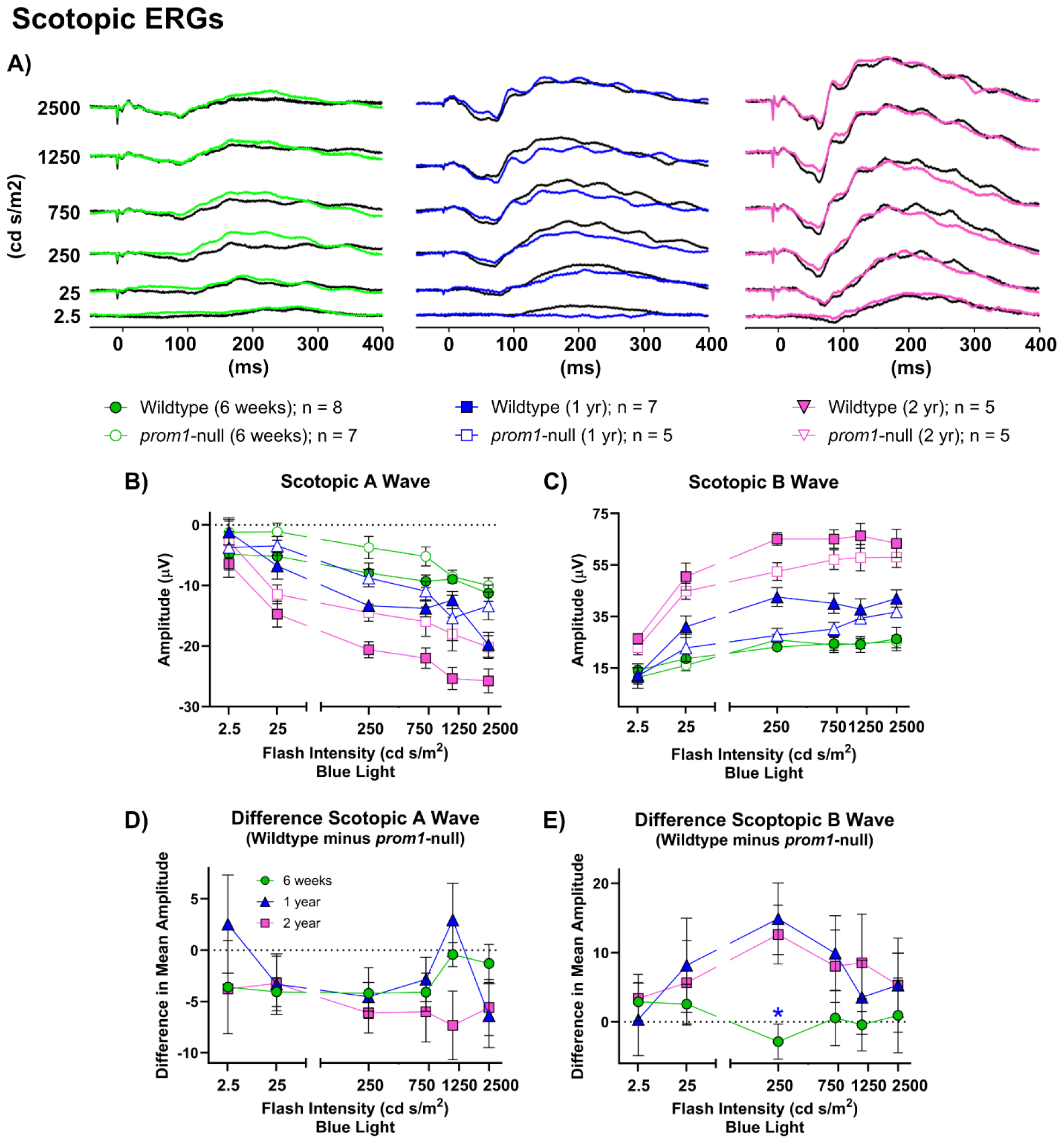
Retinal function as measured by gross scotopic ERG response amplitude and the difference between wildtype and *prom1*-null animal response amplitude across three different ages: 6 weeks (green circles), 1 year (blue triangles), and 2 years (pink squares). A) Waterfall plots of scotopic ERGs for 6 weeks, 1 year, and 2-year-old frogs. Wildtype traces are black while the mutant traces are colored. B) Scotopic A-wave response amplitude from wildtype and *prom1*-null animals. C) Scotopic B-Wave response from wildtype and *prom1*-null animals. D) The difference in A-wave response amplitude between wildtype and *prom1*-null animals. E) The difference in scotopic B-wave response amplitude between wildtype and *prom1*-null animals. Number of animals and respective symbols are denoted under the waterfall plots. Difference data were calculated and graphed as the mean difference of wildtype minus *prom1*-null animals ± SEM. Data were analyzed using Two-Way ANOVA to segregate effects of age and light intensity. For differences data (D,E), statistical significance is denoted above the group that is different, with colours of asterisks indicating the statistical significance for the comparison group. *Asterisks:* * p < 0.05. Data from 6 week old animals were published previously(Carr et al., 2021).

**Figure 7.**
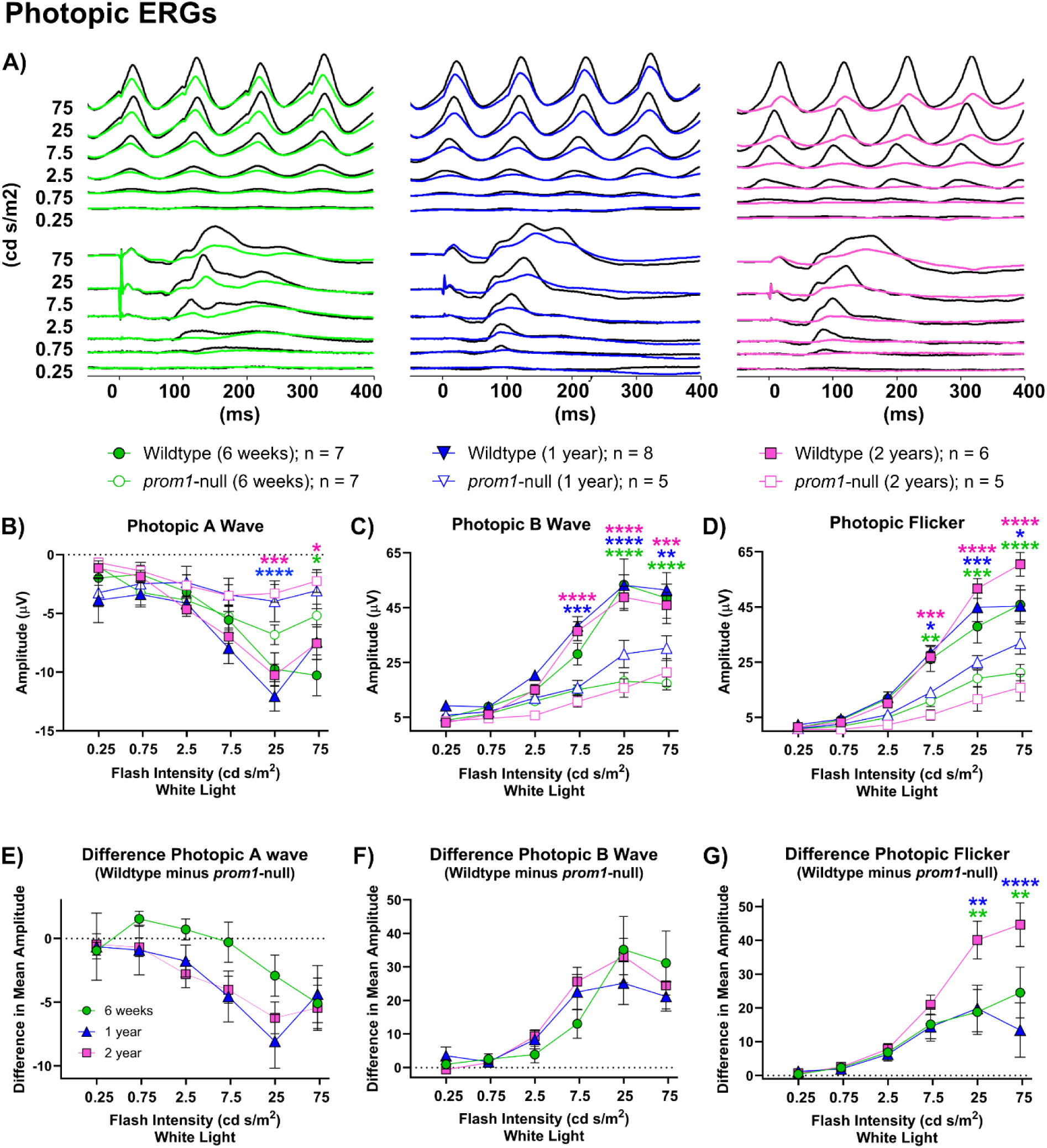
Retinal function as measured by gross photopic ERG response and the difference between wildtype and *prom1*-null animal response amplitude across three different ages: 6 weeks (green circles), 1 year (blue triangles), and 2 years (pink squares). A) Waterfall plots of photopic ERG and flicker responses for 6 week, 1 year, and 2 year old frogs. Wildtype traces are black while the mutant traces are colored. B) Gross photopic A-wave response amplitude of wildtype and *prom1*-null animals. C) Gross photopic B-wave response amplitude of wildtype and *prom1*-null animals. D) Gross photopic flicker response amplitude of wildtype and *prom1*-null animals. E) The difference in photopic A-wave response between wildtype and *prom1*-null animals. F) The difference in photopic B-wave response amplitude between wildtype and *prom1*-null animals. G) The difference in photopic flicker response amplitude between wildtype and *prom1*-null animal. Number of animals and respective symbols are denoted under the waterfall plots. Difference data were calculated and graphed as the mean difference of wildtype minus *prom1*-null animals ± SEM. For gross photopic amplitude response (B,C,D), statistical differences on the graphs denote differences between wildtype and *prom1*-null animals for the same age group; color indicates the age. For differences data (E,F,G), statistical significance is denoted above the group that is different, with colours of asterisks indicating the statistical significance for the comparison group. *Asterisks:* * p < 0.05, ** p < 0.01, *** p < 0.001, **** p < 0.0001. Data from 6 week old animals were published previously(Carr et al., 2021).

Decreased CTBP2 immunoreactivity was evident by 6 months post-fertilization, before other obvious signs of retinal degeneration such as photoreceptor outer segment loss, and was correlated with an increase in size and number of disorganized photoreceptor ribbon synapses in the cone pedicles of aging frogs as observed with TEM in 8 month old frogs (Fig. 8A; Supplementary Fig. S6). As animals aged greater than 2 years (3-6 years), worsening of outer segment disorganization and the presence of very large SDD-like deposits occurred concurrently with decreased calbindin immunoreactivity in the cone inner segments (Fig. 8B). Cone opsin immunopositivity was also decreased, though some scattered cone opsin positive material remained adhered to the rod outer segments. Finally, severely degenerated and disorganized retinas exhibited loss of calbindin immunoreactivity in the cone inner segments and a subset of bipolar cells, accompanied by thinning of the inner plexiform layer (Fig. 8C).

**Figure 8.**
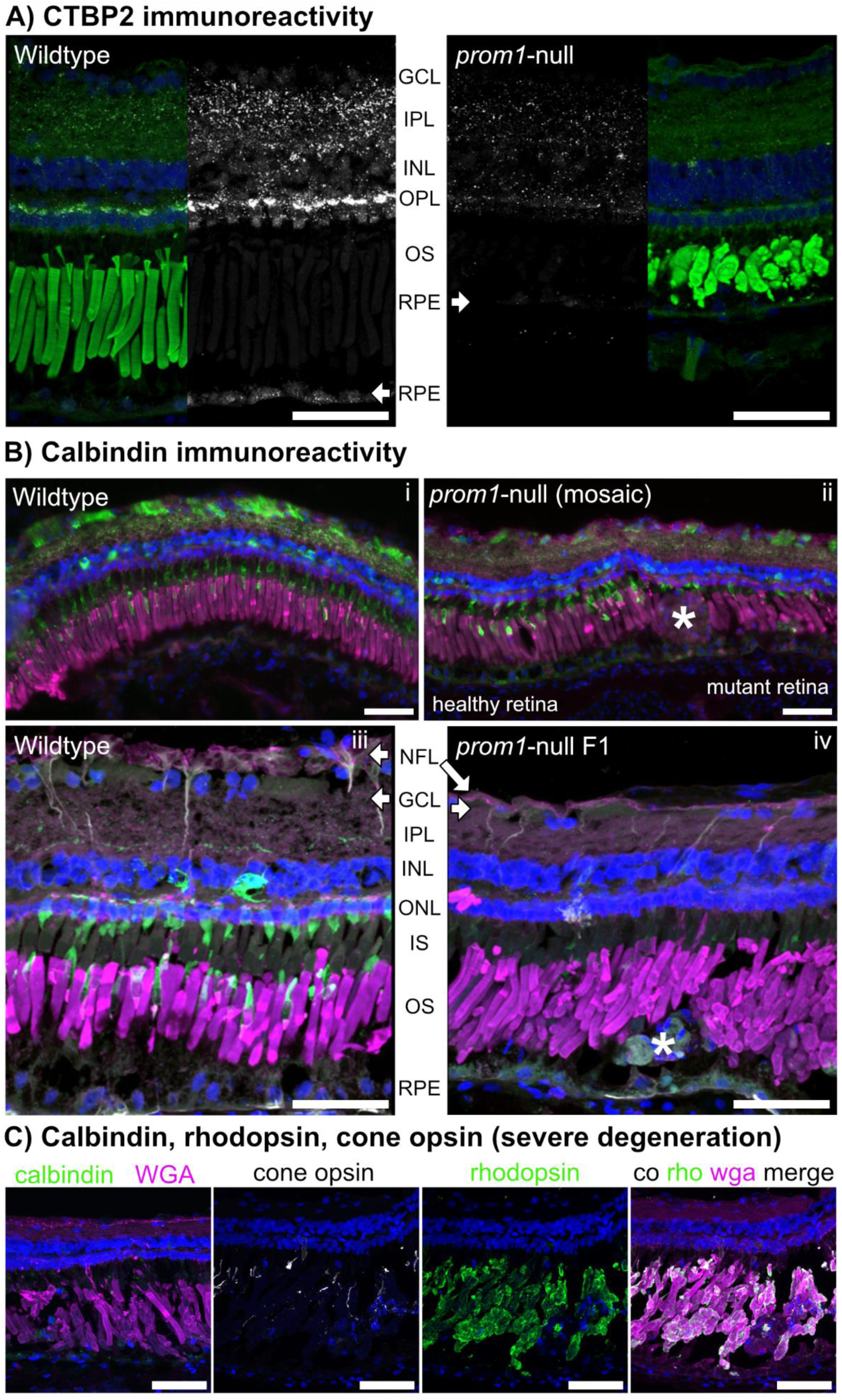
*Prom1*-null mutations result in a cone-rod dystrophy type retinal degeneration marked by changes in immunoreactivity to CTBP2, calbindin, and cone opsin. A) There is significant loss of CTBP2 immunoreactivity in the central retinas and RPE of *prom1*-null frogs aged 6 months. B) In animals aged ≥ 2 years, there is significant loss of calbindin immunoreactivity that starts off as just loss in the cone outer segments with calbindin labelling in healthy retina on the left, but loss of calbindin in mutant retina with SDD-like deposit (*) on the right (ii; mosaic F0 animal). This loss of calbindin expands to immunoreactivity in a subset of bipolar cells concurrent with inner plexiform and nerve fibre layer thinning (iv, F1 animal). There is also increased vimentin labeling in cellular debris deposits (iv, grey; *). C) Severely degenerated retinas lose calbindin immunoreactivity completely and have significant thinning of the inner plexiform layer. Some cone opsin immunoreactivity remains within the outer segment layer, but it is greatly reduced. Rhodopsin and wheat germ agglutinin immunoreactivity are preserved. i,ii are taken with the Leica DM600B epifluorescence microscope. *Scale bars* = 50 µm. *Abbreviations:* NFL, nerve fibre layer; GCL, ganglion cell layer; IPL, inner plexiform layer; INL, inner nuclear layer; OPL, outer plexiform layer; ONL, outer nuclear layer; IS, inner segments; OS, outer segments; RPE, retinal pigment epithelium. *Labels:* (A): green = wheat germ agglutinin, blue = Hoechst, white = CTBP2. (B): green = calbindin (cone inner segments and bipolar cells), blue = Hoechst, magenta = wheat germ agglutinin, white = vimentin. *Number of Animals:* Wildtype, n = 4; *prom1*-null, n = 5.

## DISCUSSION

Here, we present evidence that *prom1*-null frogs develop SDD-like deposits of cellular debris in the outer segment layer and a cone-rod dystrophy phenotype. We also present data that support the hypothesis that retinal degeneration caused by *prom1*-null mutations is not due to direct effects on photoreceptor morphogenesis. It is instead a slow process associated with build up of cellular debris in the outer segment layer and RPE atrophy, which is likely driven by RPE dysfunction and death prior to photoreceptor degeneration. These data suggest a novel mechanism of retinal degeneration for *prom1*-null mutations and demonstrate that *prom1*-null frogs may also be a suitable model organism for the study of dry AMD-related hallmarks such as deposit formation, metabolic waste build up within the RPE, RPE atrophy, and non-neovascular retinal atrophy.

Loss of prom1 results in outer segment membrane overgrowth and loss of structural integrity (Fig. 1), but not loss of membrane evaginations or inhibition of outer segment morphogenesis, as evidenced by long-term survival of the dysmorphic rods and cones reported here. Cones are more severely affected than rods (Carr et al., 2021) and *prom1*-null frogs eventually develop a cone-rod dystrophy phenotype evidenced by loss of calbindin immunoreactivity (Fig. 10B), thinning of the inner plexiform layer (Fig. 8C), and progressive decreases in photopic ERG flicker function (Fig. 7), whereas rod outer segments express rhodopsin (Fig. 8C) and scotopic ERG response remains intact (Fig. 6). We also detected changes in ribbon synapse immunoreactivity (Fig. 8A), size, and morphology in *prom1*-null frogs with moderate to severe disease (Supplementary Fig. S6). Similar findings are present in other animal models undergoing retinal remodelling in response to severe retinal degenerative disease such as retinitis pigmentosa, indicating a common endpoint of various classes of retinal degeneration (Jones and Marc, 2005). Similar to a previous report of *prom1*-null associated retinal degeneration in mouse (Zacchigna et al., 2009), we saw no effect on vitreal blood vessel morphology or indicators of choroidal neovascularization. There were no morphological defects observed in the vitreal blood vessel structure by DOPU OCT (Fig. 4) and there were no leaky vessels as assayed by fluorescein angiography (Supplementary Fig. S7). In some older animals, choroidal vessels were faintly visible through the RPE when using angiography, but this was also observed in wildtype animals (Supplementary Fig. S7). The frog retina is avascular, so vessels seen using fluorescein angiography are primarily the vitreal blood vessels (Miodoński and Bär, 1987). This phenotype is consistent with the autosomal recessive slow progressive retinal degeneration seen in human *PROM1*-null patients, who can take 15-50 years to develop a severe clinical phenotype (Gurudev et al., 2013). Extrapolating to the frog lifespan, which can be 15-20 years in captivity, aging of 2-4 years before significant retinal degeneration is analogous to the human clinical timeline and phenotype (Gurudev et al., 2013; Yang et al., 2008).

In addition to a cone-rod dystrophy phenotype, SDD-like deposits were present in aging *prom1*-null frogs and were visible in histology (Figs. 1-3), color fundus photography (Fig. 4), and as hyper-reflective foci using infrared and spectral domain OCT (Figs. 4). Hyper-reflective foci are becoming an increasingly recognized biomarker for increased risk of severe retinal degenerative disease, but the definition of “hyper-reflective foci” can refer to structures in the outer segment layer, the choroid, or cystoid voids within the retina (Fragiotta et al., 2021). There are many hypotheses regarding the sources of material in the lesions, including macrophages, microglia, undigested lipid or protein material, retinal exudate, or migrating RPE cells (Fragiotta et al., 2021). The deposits we observed in *prom1*-null frogs were localized to the outer segment layer, large (>30 µm), not associated with retinal exudates, and did not exhibit OCT shadowing (Fig. 4). This categorizes them as similar to hyper-reflective foci associated with severe outcomes of human retinal degenerative diseases such as retinitis pigmentosa (Kuroda et al., 2014), intermediate to severe age-related macular degeneration (Curcio et al., 2017), and Stargardt disease (Battaglia Parodi et al., 2019; Piri et al., 2015; Ritter et al., 2013). A limitation to the current study was that the OCT beam does not penetrate into the frog choroid as it does in mammalian choroid (Gong et al., 2015), making it difficult to ascertain whether *prom1*-null frogs also have significant hyper-reflective foci in this layer. Notably, increasing size and number of hyper-reflective foci are considered a precursor of geographic atrophy (Christenbury et al., 2013; Leuschen et al., 2013; Schuman et al., 2009). Morphological features of progression to geographic atrophy include reduced retinal thickness, ONL thinning, and migration of hyper-reflective foci to the inner retina (Balaratnasingam et al., 2016; Ouyang et al., 2013).

Involvement of RPE dysfunction prior to photoreceptor degeneration in our *prom1*-null frogs is supported by several lines of evidence. Firstly, there was no progressive loss of ERG response from the rods and very little progressive loss of ERG response from cones of *prom1*-null frogs between the ages of 6 weeks and two years (Figs. 6,7). Outer segment membranes are synthesized continuously throughout life (Anderson et al., 1978; Young, 1967), and complete turnover of the outer segment membranes in *Xenopus laevis* raised under normal lighting and temperature conditions should happen within approximately 28 days (Kinney and Fisher, 1978). Thus, if *prom1*-null mutations inhibited plasma membrane evaginations or some other process involved in outer segment membrane renewal, we would have expected to see significant loss of outer segments and loss of visual function on a significantly faster timescale (weeks to months). Instead, we observed significantly dysmorphic outer segments, indicating a role of prom1 protein in regulating membrane growth, but no loss of rod outer segments for ≥ 2 years.

The second line of evidence that retinal degeneration caused by *prom1*-null mutations is time-or age-dependent and involves RPE is that visual impairment and retinal degeneration in *prom1*-null frogs was preceded by ever increasing size and number of deposits of cellular debris in the subretinal space that then extended further into the outer segment layer and eventually through the ELM into the inner retina (Fig. 1-4). The deposits exhibited autofluoresce at significantly different wavelengths than the surrounding outer segment material, indicating a different source or significant protein and lipid degradation within the deposits. These deposits resembled SDD in location, appearance in color fundus photography and OCT, and reaction to the histological stains Oil Red O and BODIPY (Fig. 2). There are few animal models reported to faithfully recapitulate SDD, and those that do require conditional double-knockout of cholesterol transporters ABCA1 and ABCG1 in macrophages (Ban et al., 2018). Frog deposits were detected in color fundus photography as bluish-white lesions scattered across the retina in a regular mosaic pattern (Fig. 4). The distribution of lesions associated with *prom1*-null mutations correlates with a previous hypothesis that SDD may be associated with a specific retinal cell type, namely rods, while soft drusen are associated with cones (Curcio et al., 2013). We were unable to detect any rhodopsin-positive immunoreactivity in the deposits. We were able to detect some cone opsin positive labeling, which was restricted to honeycomb-like cell borders within the deposits, and not uniform throughout the deposit (Fig. 3). Frogs do not have a macula or visual streak, and their retinal neurons are regularly interspersed in similarly uniform mosaic pattern (Wilhelm and Gábriel, 1999). Thus, while we could not confirm directly that the frog SDD-like deposits were associated specifically with rods or cones, the consistent mosaic distribution of the deposits does indicate specific associations with retinal cell types or circuits.

Taking into consideration the localization, OCT reflectivity, and histology of the observed *prom1*-null frog SDD-like deposits, it seems likely that the cellular source of these deposits is the RPE. Evidence for this includes significant infiltration of RPE pigment granules into the deposits (Fig. 2), disordered and strong RPE65 positive immunolabelling detected around “displaced nuclei” (Fig. 3) that could represent early stages in deposit formation, build-up of membranous and vacuolated material in and around the RPE as observed in TEM images (Fig. 5), and window defects that represent RPE and choroidal pigment loss in some *prom1*-null F1 animals (Fig. 4). These window defects were best detected through the use of DOPU OCT, which shows clearly the breakdown of highly polarizing signals in the pigmented RPE and choroid (Fig. 4). 3D reconstruction of DOPU signal in large deposits also showed a core of highly depolarizing signal surrounded by non-polarizing debris (Fig. 4). This could represent a core of dead RPE material made up of depolarizing pigment granules that continues to accumulate further debris as the animal ages. Finally, retina/RPE of 2-year-old *prom1*-null animals are, on average, thinner than WT retinas as measured by OCT (Supplementary Fig. S4).

RPE are terminally differentiated cells that are never replaced throughout the lifetime of the organism (Strauss, 2005). One of the primary functions of the RPE is to phagocytose shed outer segment membranes, creating a lifetime of significant oxidative stress as the lipid membranes are metabolized (Strauss, 2005). Oxidative stress can cause a process of cellular dysfunction and de-differentiation termed epithelial mesenchymal transition (EMT) (Inumaru et al., 2009). EMT occurs when differentiated and polarized epithelial cells, such as the RPE, begin to lose their epithelial phenotype (polarized, adhesive, avascular) and begin to adopt a mesenchymal phenotype (non-polar, migratory, potentiating/self-renewing) (Inumaru et al., 2009). Some studies have reported that RPE cells may undergo EMT during severe retinal degenerative disease such as age-related macular degeneration (Datta et al., 2017). Characteristics associated with this change are increased migratory activity, invasiveness into surrounding tissues, elevated resistance to apoptosis, and increased production of extracellular matrix and cytoskeletal components such as vimentin, N-cadherin, β-catenin, and fibronectin (Kalluri and Weinberg, 2009). We showed that there is a similar pattern of relative increased vimentin in the large deposits in *prom1*-null frogs and in the RPE surrounding these deposits (Fig. 3C).

We propose that loss of *prom1* results in RPE death though a mechanism that involves increased phagocytic and oxidative stress caused by the dysmorphic *prom1*-null photoreceptor outer segments. As shown here, and in our previous work (Carr et al., 2021), *prom1*-null outer segments lack structural integrity, especially the cones, and this loss of integrity could impact outer segment shedding, leading to an increased amount of material for the RPE to phagocytose and process (Supplementary Fig. S5). Increased phagocytic load could cause oxidative stress and increased build-up of reactive oxygen and nitrogen species and indigestible material within the RPE. Increased oxidative stress could then lead to EMT, causing even greater RPE dysfunction and creating a positive feedback loop of stress, damage, and eventual RPE death. Effects corresponding to this were observed in the increased 4 channel RPE autofluorescence (Fig. 1L), increased red autofluorescence (Fig. 2C,D), increased vimentin expression in the SDD-like deposits (Fig. 3) and in the TEM images of the RPE that showed particles of undigested membrane and vacuole materials ≥1 µm (Fig. 5). This proposed mechanism is similar to age-related build-up of lipofuscins and other materials in the RPE, which can also lead to decreased autophagic activity and RPE atrophy (Brunk and Terman, 2002). Accumulation of fluorescent metabolic byproducts of the visual transduction cycle and outer segment membrane phagocytosis is associated with photoreceptor degeneration and RPE death in Stargardt disease and atrophic (dry) age-related macular degeneration (AMD). This model also corresponds to the clinical manifestation of PROM1-associated retinal disease in humans, which is often diagnosed as cone-rod dystrophy, retinitis pigmentosa with cone involvement, or juvenile bull’s eye macular degeneration (Permanyer et al., 2010; Pras et al., 2009; Zhang et al., 2007). Autosomal dominant *PROM1* disease (R373C) is often misdiagnosed as Stargardt disease before genetic testing occurs (Michaelides et al., 2010; Yang et al., 2008).

In summary, we show here that retinal degeneration in *prom1*-null frogs is slow and has a cone-rod dystrophy phenotype associated with SDD-like deposit formation and RPE atrophy. On the basis of these data, we hypothesize that the mechanism for recessive *prom1*-null associated blindness is not direct effects on photoreceptor morphogenesis, but instead likely involves dysfunction and death of the RPE that precedes photoreceptor degeneration. Finally, our *prom1*-null frogs showed analogous effects to some key hallmarks of dry AMD, including the presence of deposits, potential dysfunction and death of the RPE, accumulation of lipofuscins and autofluorescent material in the RPE, and non-neovascular retinal atrophy. Thus, further study and characterization of this model could also lead to significant findings relevant to AMD in addition to inherited blindness.

## ACKNOWLEDGMENTS

Research funded by the Canadian Institutes for Health Research (OLM; PJT-155937, PJT-156072), the Natural Sciences and Engineering Research Council of Canada (OLM; RGPIN-2015-04326), the Edwina and Paul Heller Memorial Fund (BJC), a Michael Smith Foundation for Health Research (MSFHR) Research Trainee Award (BJC), a BrightFocus Foundation Macular Degeneration Postdoctoral Award (BJC; M2021001F), and University of Alberta Startup Funds (BJC). Acknowledgement is made to the donors of Macular Degeneration Research, a program of the BrightFocus Foundation, for support of this research. Transmission electron microscopy sample preparation and imaging supported by Dr. Kacie Norton at the Advanced Microscopy Facility (University of Alberta, Dept. of Biological Sciences) and Dr. Sara Amidian at the Cell Imaging Facility (University of Alberta, Cross Cancer Institute). Purchase of the Zeiss LSM 800 and LSM 880 with Airyscan and the Phoenix Micron IV funded by an infrastructure grant from the Canadian Foundation for Innovation (OLM; University of British Columbia). RPC1 rabbit ultrastructural volumes (Supplementary Fig. S6) were accessed using the Viking Database (Anderson et al., 2011; Pfeiffer et al., 2020), funded by NIH EY028927 (Bryan W. Jones). Prom1 immunolabeling signal intensity was measured in FIJI using a macro created by Dr. Elizabeth Kugler (EKugler_11082021.ijm; Zeeks – Art for Geeks Ltd).

## CONTRIBUTIONS

BJC designed, performed, and analyzed all experiments in this manuscript as a postdoctoral fellow, with the exception of the DOPU OCT imaging. BJC created all figures and wrote the manuscript. DS performed the DOPU OCT imaging in *prom1*-null and wildtype frogs, performed preliminary image analysis, and helped to write the methods section for DOPU OCT. JS and MJJ built the DOPU OCT system and wrote the methods section for DOPU OCT. XL performed preliminary analysis of retinal thickness calculated according to OCT (Supplementary Fig. S4). BJC, OLM and MJJ are the principal investigators of the labs where these studies and data analysis were carried out.

## SUPPLEMENTARY FIGURES

**Supplementary Figure S1.**
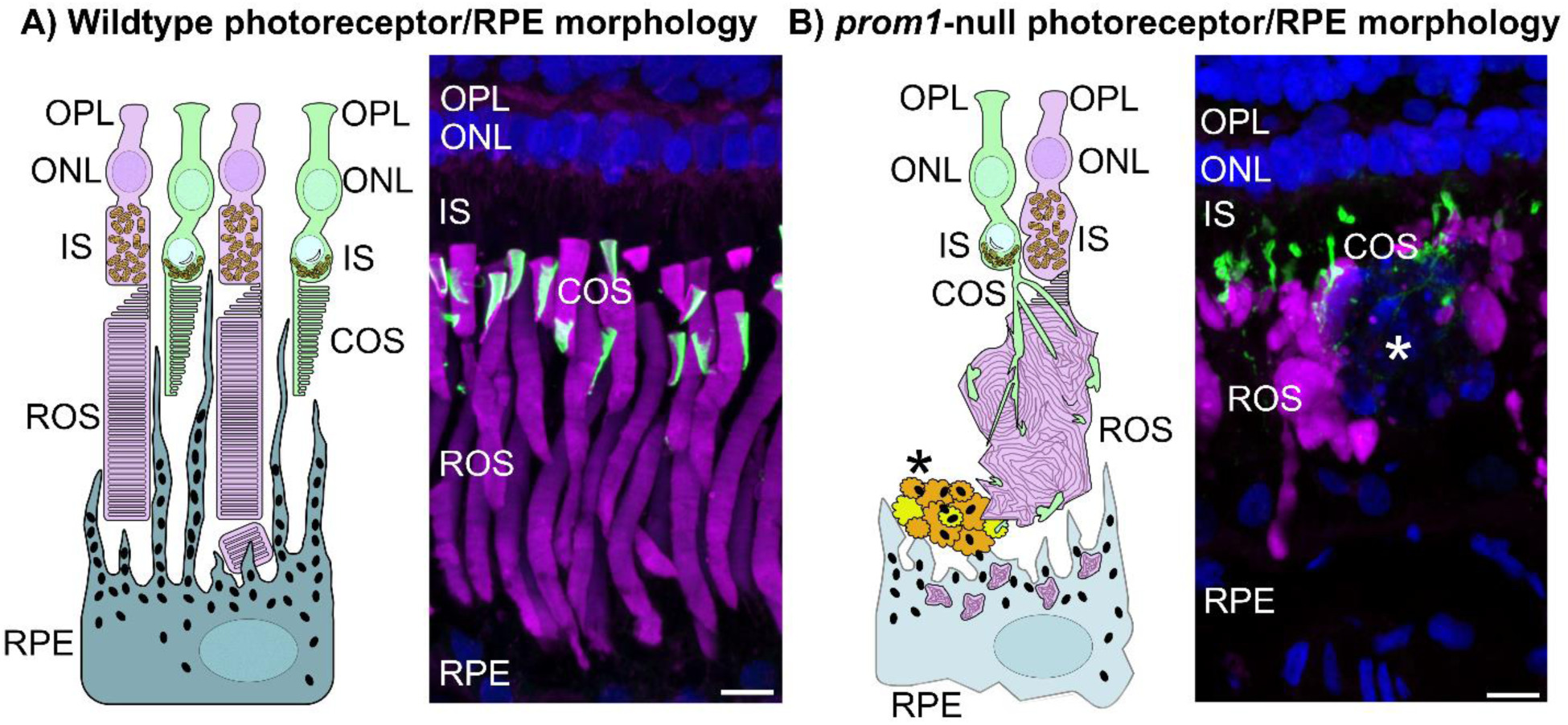
Schematic of photoreceptor/RPE morphology in wildtype (A) and *prom1*-null (B) frogs. A) Wildtype outer segment morphology is highly organized in both rods and cones. Rods (magenta) comprise discrete discs of membrane whereas cones (green) comprise layered lamellae. B) Outer segment morphology in *prom1*-null frogs is significantly dysmorphic. Rod outer segments are bulbous, shortened, and comprise overgrown whorls of membrane within the plasma membrane. Cone outer segments are stretched and fragmented, and cling to the outside of rod outer segments. There is also the presence of heterogeneous deposits of debris in the outer segment layer of *prom1*-null frogs (*), that stains with Hoechst 33342 nuclear stain, but not for markers for outer segment material (WGA, rhodopsin, cone opsin). Scale bars = 10 µm. *Micrograph labels:* Magenta, WGA; Green, cone opsin; blue, Hoechst 33342 nuclear stain. *Abbreviations:* RPE, retinal pigment epithelium; ROS, rod outer segment; IS, inner segment; ONL, outer nuclear layer; OPL, outer plexiform layer; COS, cone outer segment; WGA, wheat germ agglutinin.

**Supplementary Figure S2.**
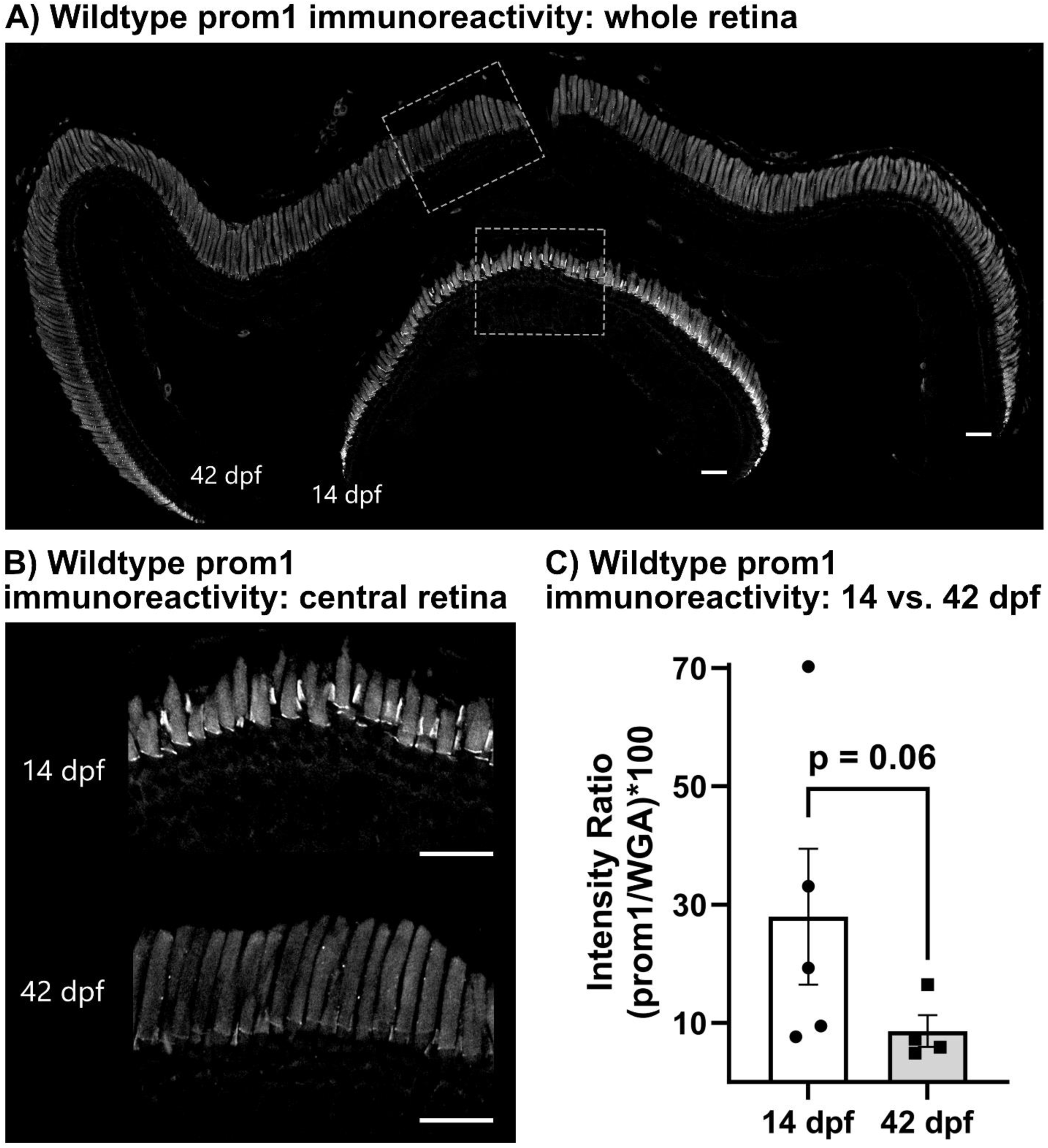
Prom1 protein immunoreactivity in wildtype animals aged 14– and 42– days post fertilization. Relative prom1 immunoreactivity decreases in the central retina as animals age and outer segments reach adult size. Scale bar = 50 µm. Bar graph demonstrates mean ± SEM. *Number of animals:* 14 dpf, n = 5; 42 dpf, n = 4.

**Supplementary Figure S3.**
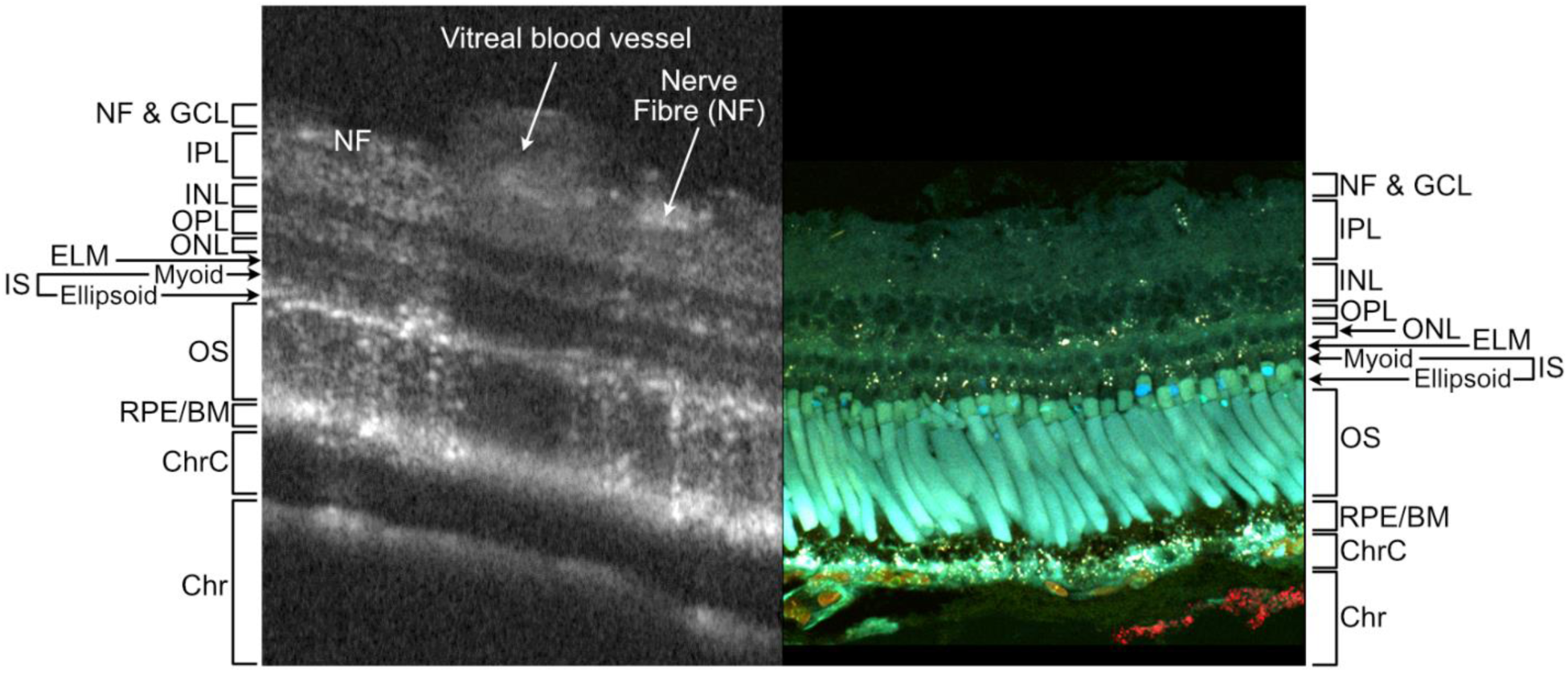
Interpretation of wildtype *Xenopus laevis* retina as imaged by OCT (left) by comparison to endogenous retinal autofluorescence (right). Retinal layers and primary structures are visible using OCT and autofluorescence. *Abbreviations:* NFL & GCL, nerve fibre layer & ganglion cell layer; IPL, inner plexiform layer; INL, inner nuclear layer; OPL, outer plexiform layer; ONL, outer nuclear layer; ELM, external limiting membrane; IS, inner segments; OS, outer segments; RPE/BM, retinal pigment epithelium & Bruch’s membrane; ChrC, choriocapillaris; Chr, choroid.

**Supplementary Figure S4.**
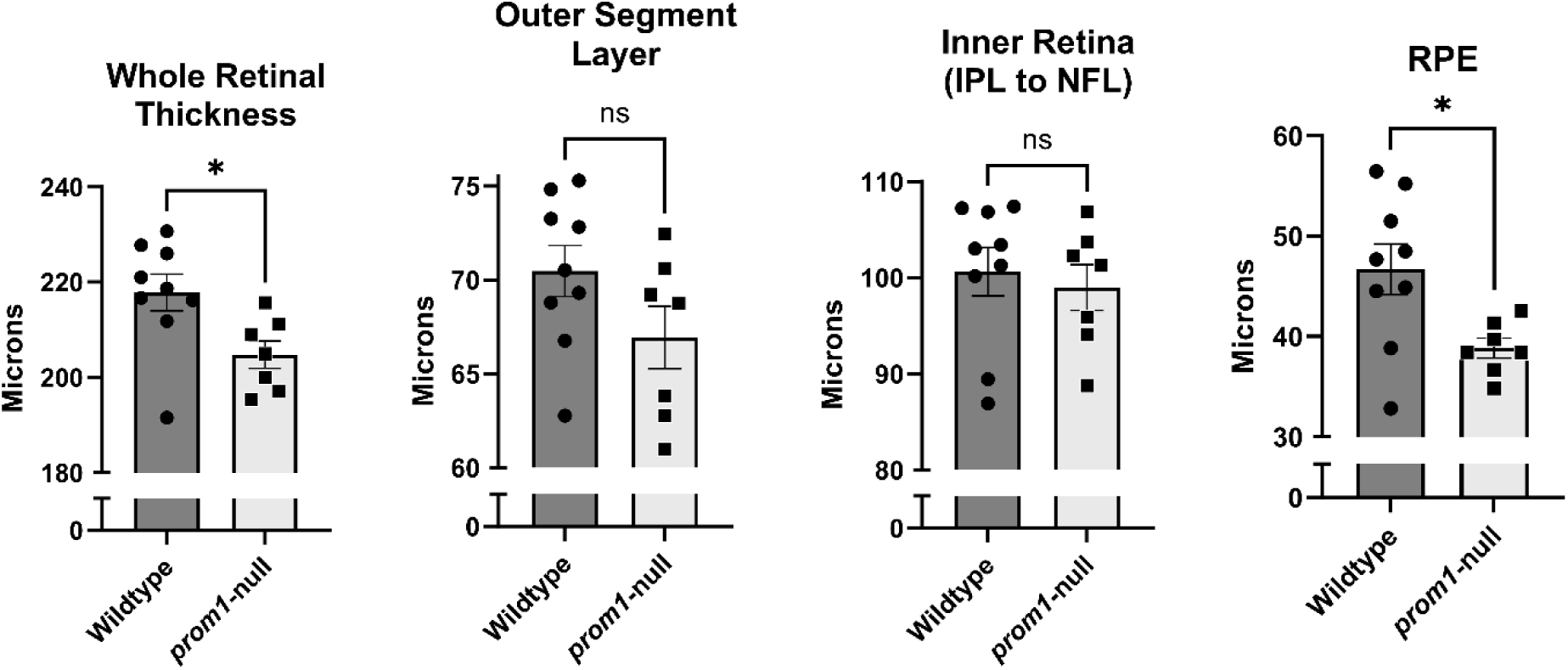
Comparison of the thickness of retinal layers in 2-year-old wildtype and F0 *prom1*-null animals. The RPE represented the greatest contributor to overall thinning of the retinal in *prom1*-null frogs. Graphs represent the means ± SEM. *Number of animals:* Wildtype, n = 9; *prom1*-null, n = 7.

**Supplementary Figure S5.**
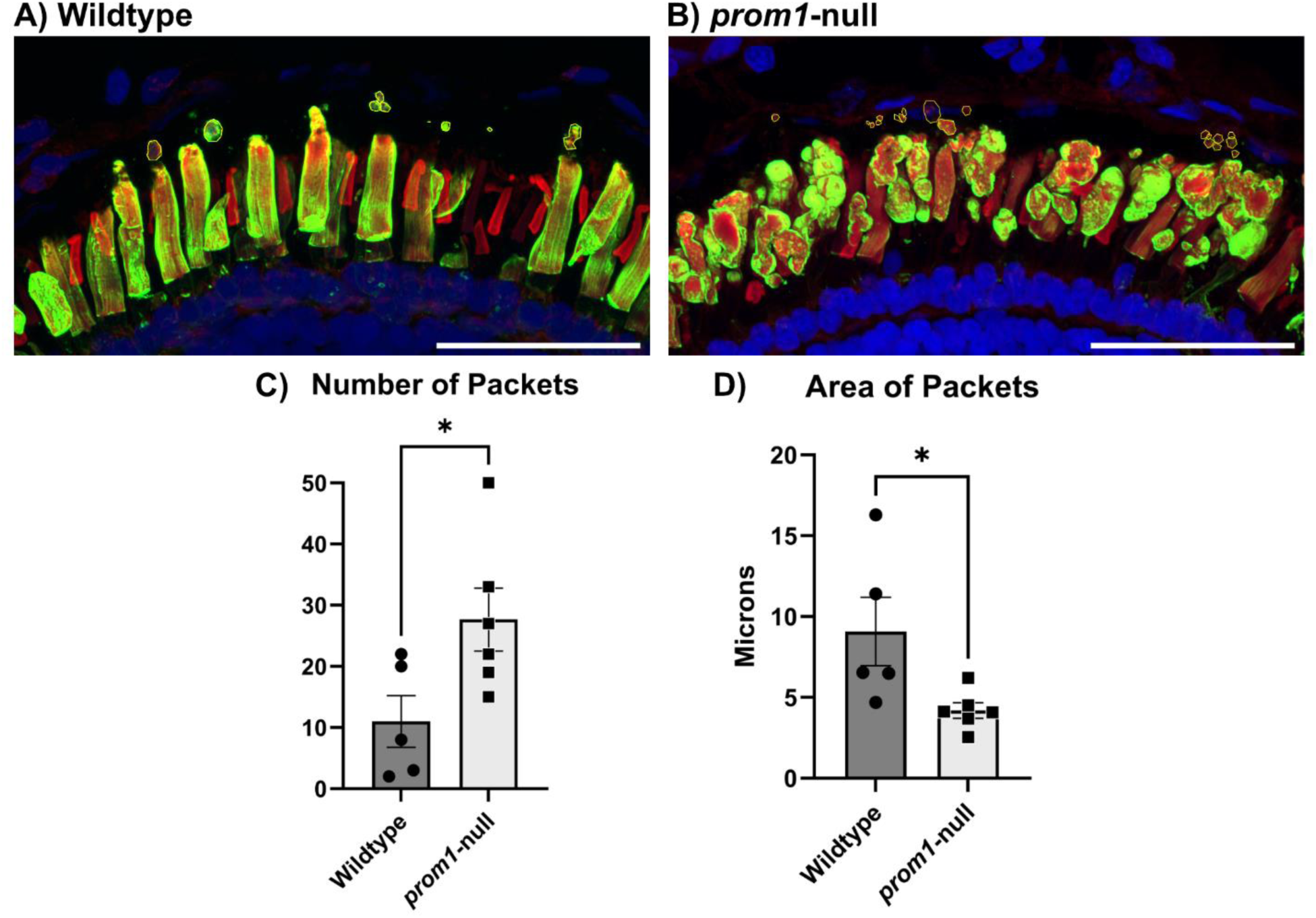
Comparison of the size and number of shed outer segment packets in wildtype versus *prom1*-null mutant tadpoles aged 14 dpf. Scale bars = 50 µm. Data are represented as mean ± SEM. *Number of animals:* wildtype, n = 5; *prom1*-null n = 6. *Statistics:* Student’s t-test, unpaired, two-tailed; * p < 0.05.

**Supplementary Figure S6.**
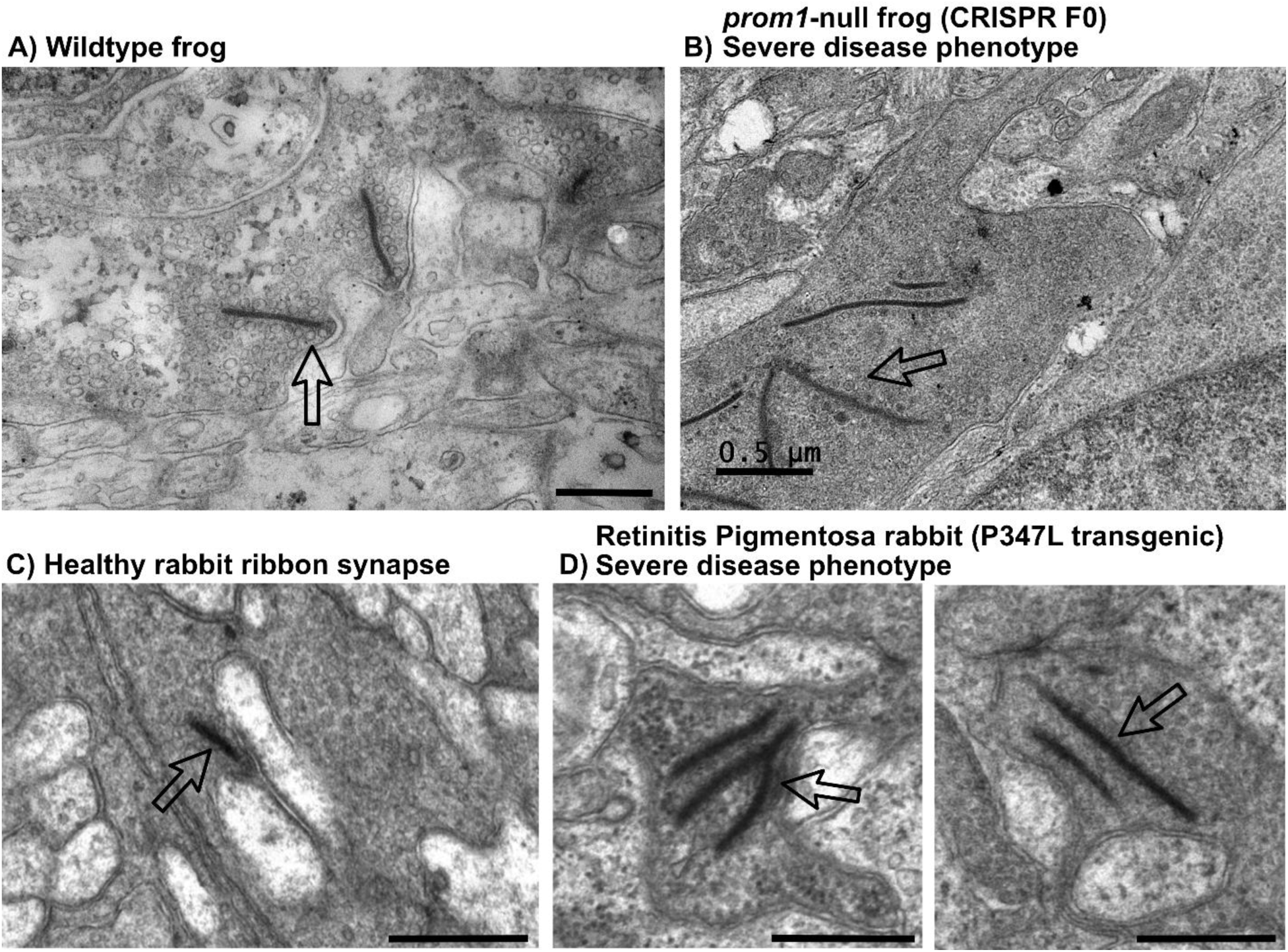
A comparison of morphology changes in photoreceptor synapses between a wildtype frog, a *prom1*-null frog, and a rabbit model of retinitis pigmentosa (P347L). The *prom1*-null frog and P347L rabbit have a severe phenotype and retinal synaptic remodelling. Remodelling features are very similar between the two animals and the two models of retinal degeneration, indicating similarities in the downstream processes of severe retinal degeneration. Scale bar = 0.5 µm.

**Supplementary Figure S7.**
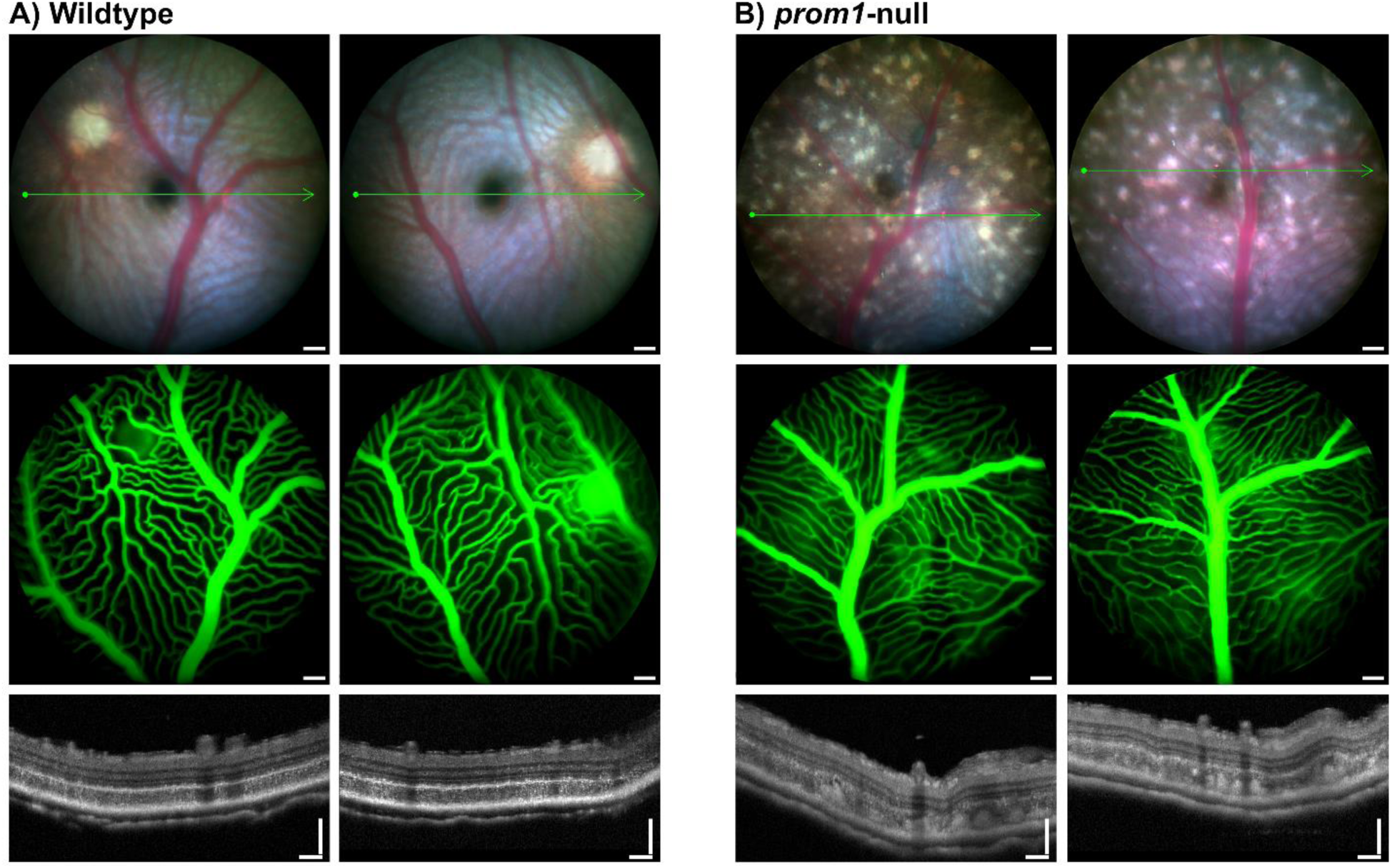
Fluorescein angiography in 3-year-old wildtype and *prom1*-null mutant frogs. There was little observed difference in the vitreal blood vessel structure and there were no leaky vessels or significant bleed through to the choroidal vessels near the large lesions. Scale bars = 100 µm. *Number of animals:* Wildtype, n = 9, *prom1*-null, n = 8.

**Supplementary Table S1.**
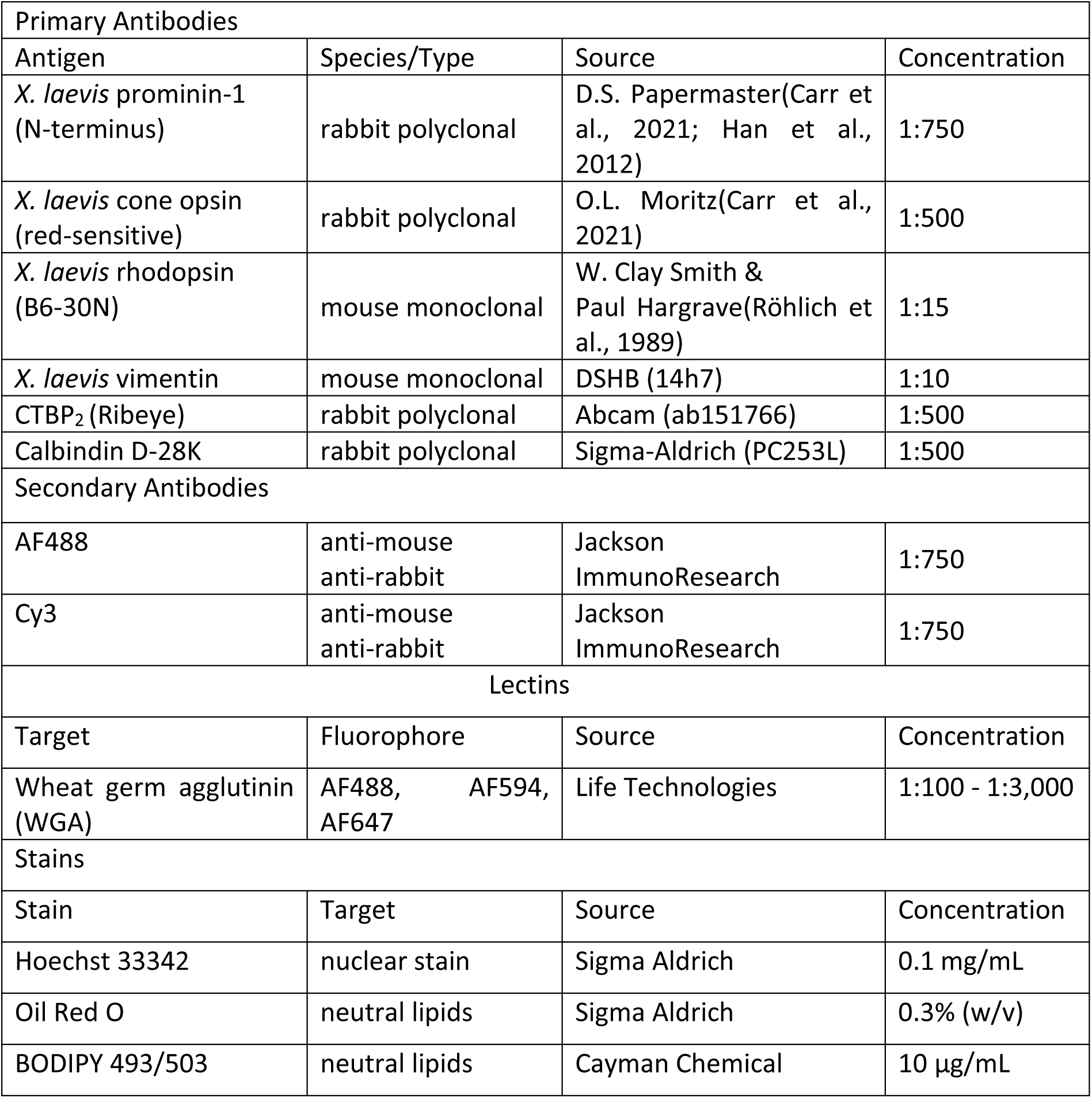
Antigens, species/type, source, and working dilutions of antibodies used in this study.

| Primary Antibodies |  |  |  |
| --- | --- | --- | --- |
| Antigen | Species/Type | Source | Concentration |
| *X. laevis* prominin-1 (N-terminus) | rabbit polyclonal | D.S. Papermaster(Carr et al., 2021; Han et al., 2012) | 1:750 |
| *X. laevis* cone opsin (red-sensitive) | rabbit polyclonal | O.L. Moritz(Carr et al., 2021) | 1:500 |
| *X. laevis* rhodopsin (B6-30N) | mouse monoclonal | W. Clay Smith & Paul Hargrave(Röhlich et al., 1989) | 1:15 |
| *X. laevis* vimentin | mouse monoclonal | DSHB (14h7) | 1:10 |
| CTBP<sub>2</sub> (Ribeye) | rabbit polyclonal | Abcam (ab151766) | 1:500 |
| Calbindin D-28K | rabbit polyclonal | Sigma-Aldrich (PC253L) | 1:500 |
| Secondary Antibodies |  |  |  |
| AF488 | anti-mouse<br>anti-rabbit | Jackson ImmunoResearch | 1:750 |
| Cy3 | anti-mouse<br>anti-rabbit | Jackson ImmunoResearch | 1:750 |
| Lectins |  |  |  |
| Target | Fluorophore | Source | Concentration |
| Wheat germ agglutinin (WGA) | AF488, AF594, AF647 | Life Technologies | 1:100 - 1:3,000 |
| Stains |  |  |  |
| Stain | Target | Source | Concentration |
| Hoechst 33342 | nuclear stain | Sigma Aldrich | 0.1 mg/mL |
| Oil Red O | neutral lipids | Sigma Aldrich | 0.3% (w/v) |
| BODIPY 493/503 | neutral lipids | Cayman Chemical | 10 µg/mL |

**Supplementary Table S2.** Statistics for photopic flash and flicker ERG. *Asterisks:* * p < 0.05, ** p < 0.01, *** p < 0.001, **** p < 0.0001.

| Photopic A-Wave | wildtype vs. *prom1* -null | | | Two-way ANOVA<br>Interaction effect: $p < 0.0034$<br>Simple main light intensity effect: $p < 0.0001$<br>Simple main genotype effect: $p < 0.0001$ |
| --- | --- | --- | --- | --- |
| Light Intensity (cd s/m<sup>2</sup>) | 6 weeks | 1 year | 2 year |  |
| 0.25 | ns | ns | ns |  |
| 0.75 | ns | ns | ns |  |
| 2.5 | ns | ns | ns |  |
| 7.5 | ns | ns | ns |  |
| 25 | ns | **** | *** |  |
| 75 | * | ns | * |  |

| Photopic B-Wave | wildtype vs. *prom1* -null | | | Two-way ANOVA<br>Interaction effect: $p < 0.0001$<br>Simple main light intensity effect: $p < 0.0001$<br>Simple main genotype effect: $p < 0.0001$ |
| --- | --- | --- | --- | --- |
| Light Intensity (cd s/m<sup>2</sup>) | 6 weeks | 1 year | 2 year |  |
| 0.25 | ns | ns | ns |  |
| 0.75 | ns | ns | ns |  |
| 2.5 | ns | ns | ns |  |
| 7.5 | ns | *** | **** |  |
| 25 | **** | **** | **** |  |
| 75 | **** | ** | *** |  |

| Photopic Flicker | wildtype vs. *prom1* -null | | | Two-way ANOVA<br>Interaction effect: $p < 0.0001$<br>Simple main light intensity effect: $p < 0.0001$<br>Simple main genotype effect: $p < 0.0001$ |
| --- | --- | --- | --- | --- |
| Light Intensity (cd s/m<sup>2</sup>) | 6 weeks | 1 year | 2 year |  |
| 0.25 | ns | ns | ns |  |
| 0.75 | ns | ns | ns |  |
| 2.5 | ns | ns | ns |  |
| 7.5 | ** | * | *** |  |
| 25 | *** | *** | **** |  |
| 75 | **** | * | *** |  |

| Difference Photopic Flicker | wildtype minus *prom1* -null | | | Two-way ANOVA<br>Interaction effect: $p = 0.0165$<br>Simple main light intensity effect: $p < 0.0001$<br>Simple main genotype effect: $p = 0.0006$ |
| --- | --- | --- | --- | --- |
| Light Intensity (cd s/m<sup>2</sup>) | 6 weeks<br>vs. 1 year | 6 weeks<br>vs. 2 year | 1 year<br>vs. 2 year |  |
| 0.25 | ns | ns | ns |  |
| 0.75 | ns | ns | ns |  |
| 2.5 | ns | ns | ns |  |
| 7.5 | ns | ns | ns |  |
| 25 | ns | ** | ** |  |
| 75 | ns | ** | **** |  |

